# *Drosophila* Neuronal Glucose 6 Phosphatase is a Modulator of Neuropeptide Release that Regulates Muscle Glycogen Stores via FMRFamide Signaling

**DOI:** 10.1101/2023.11.28.568950

**Authors:** Tetsuya Miyamoto, Sheida Hedjazi, Chika Miyamoto, Hubert Amrein

**Affiliations:** Department of Cell Biology and Genetics School of Medicine Texas A&M University Bryan, Texas

## Abstract

Neuropeptides (NPs) and their cognate receptors are critical effectors of diverse physiological processes and behaviors. We recently reported of a non-canonical function of the *Drosophila Glucose-6-Phosphatase* (*G6P*) gene in a subset of neurosecretory cells in the CNS that governs systemic glucose homeostasis in food deprived flies. Here, we show that *G6P* expressing neurons define 6 groups of neuropeptide secreting cells, 4 in the brain and 2 in the thoracic ganglion. Using the glucose homeostasis phenotype as a screening tool, we find that neurons located in the thoracic ganglion expressing FMRFamide neuropeptides (*FMRFa^G6P^* neurons) are necessary and sufficient to maintain systemic glucose homeostasis in starved flies. We further show that *G6P* is essential in *FMRFa^G6P^* neurons for attaining a prominent Golgi apparatus and secreting neuropeptides efficiently. Finally, we establish that *G6P* dependent FMRFa signaling is essential for the build-up of glycogen stores in the jump muscle which expresses the receptor for FMRFamides. We propose a general model in which the main role of *G6P* is to counteract glycolysis in peptidergic neurons for the purpose of optimizing the intracellular environment best suited for the expansion of the Golgi apparatus, boosting release of neuropeptides and enhancing signaling to respective target tissues expressing cognate receptors.

**SIGNIFICANCE STATEMENT:** Glucose-6-phosphtase (G6P) is a critical enzyme in sugar synthesis and catalyzes the final step in glucose production. In *Drosophila* - and insects in general - where trehalose is the circulating sugar and Trehalose phosphate synthase, and not G6P, is used for sugar production, G6P has adopted a novel and unique role in peptidergic neurons in the CNS. Interestingly, flies lacking *G6P* show diminished Neuropeptide secretions and have a smaller Golgi apparatus in peptidergic neurons. It is hypothesized that the role of G6P is to counteract glycolysis, thereby creating a cellular environment that is more amenable to efficient neuropeptide secretion.

## INTRODUCTION

Neuropeptides (NPs) play central roles in modulating physiology and behaviors. In humans, more than one hundred NPs have been identified, and most of them act through known G protein coupled receptors expressed in specific neurons of the brain or target cells in non-neuronal organs and tissues (Nässel and Zandawala, 2019; van den Pol, 2012). *Drosophila melanogaster* has become an important model system for investigating the diverse functions of NPs and their receptors, due to the expansive array of molecular genetic tools, the ability to visualize electrophysiological activity and the availability of powerful behavioral assays. More than 40 NP genes have been described in the fruit fly, many of which encode multiple peptides, and for many of these, cognate receptors have been identified (Nässel and Zandawala, 2019). We recently reported expression of the gluconeogenic enzyme Glucose-6-phosphatase (G6P) in small subpopulations of peptidergic neurons in the fly CNS. By expressing a CFP/YFP based glucose sensor, we showed that G6P is functional in these cells for not only maintaining intracellular glucose levels when flies are food deprived, but also as a critically important enzyme necessary for the conversion of alanine to glucose (Miyamoto and Amrein, 2019). At the systemic level, we found that starved *G6P* mutant flies failed to maintain whole body glucose homeostasis, a phenotype that is not only rescued by expression of a *G6P* transgene, but also by activation of *G6P* neurons using the heat-activated channel TRPA1 in *G6P* mutant flies. Taken together, these observations indicated that the role of *G6P* in these neurons is not to produce glucose, but to facilitate peptidergic signaling, disruption of which causes loss of whole-body glucose homeostasis in starved flies.

In mammals, G6P is mainly known as a key liver and kidney enzyme during gluconeogenesis and glycogenolysis, metabolic processes upregulated in food-deprived animals. G6P hydrolyses g-6-p to glucose, which is then released into the blood stream to maintain glucose homeostasis. These two pathways are highly conserved across different animal lineages including insects, where they take place in the adipose (fat body) and Malpighian tubules, tissues analogous to the mammalian liver and kidney, respectively. However, the last enzymatic step is distinct in insects, whereby g-6-p is converted to trehalose, the main circulating insect sugar, using the enzyme trehalose 6 phosphate synthase 1 (TPS1) to ensure sugar homeostasis when flies are starved (Matsuda et al., 2015; Thompson, 2003). Thus, the existence of a conserved *Drosophila* G6P enzyme in general, and its expression in peptidergic neurons specifically, which are not known to be involved in systemic sugar homeostasis, raises intriguing questions about its function in NP signaling. First and foremost, what is the neuronal identity of the approximately 30 *G6P-GAL4* expressing neurons in the CNS, beyond the fact that they express DIMMED, a marker for large NP producing neurons (Hewes et al., 2003)? Second, which of these neurons mediate whole body glucose homeostasis in starved flies, the only know function of *G6P* in insects? Third, what is the cellular role of G6P in peptidergic neurons? And fourth, what is the primary systemic phenotype that ultimately leads to failed glucose homeostasis in starved *G6P* mutant flies?

In this paper, we provide answers to these questions. We determined the identity of *G6P* neurons in the fly CNS by correlating *G6P-GAL4* expression with the location of NP producing neurons. Specifically, we identified subsets of large neuropeptide F (NPF), Orcokinin A (OrcoA), pigment-dispersing factor (Pdf), FMRFamide and IPNamide - positive neurons that also express *G6P-GAL4.* By inhibiting neural activity and rescuing G6P function in *G6P* mutant flies in each of these subsets, we show that *FMRFa* expressing *G6P* neurons (*FMRFa^G6P^*) located in the thoracic ganglia alone are necessary and sufficient to maintain whole body glucose homeostasis in food deprived flies. We further narrowed down the precise subset of *FMRFa^G6P^* neurons using the split GAL4 system and found that they correspond to four pairs of ventrally located large FMRFa expressing neurons (T1v, T2v and T3v and T2va neurons; Schneider et al., 1993b). Moreover, we have uncovered the first cellular phenotypes associated with *FMRFa^G6P^* neurons, namely a smaller Golgi apparatus and reduced NP secretion from *FMRFa^G6P^* neurons. Lastly, we describe the primary systemic role mediated by *G6P* and *FMRFa*, by homing in on the FMRFa receptor (FMRFaR). Specifically, we show that *FMRFaR* is expressed in the jump muscle and that muscle glycogen content of *G6P^-/-^*, *FMRFa^-/-^* and *FMRFaR^-/-^* flies is reduced by about 50 percent, compared to wild type controls. These observations indicate that FMRFa-FMRFaR signaling is essential to build up glycogen stores under normal feeding conditions, and they suggest that the loss of whole-body glucose homeostasis observed of starved *G6P* mutants is due to the inability of these flies to draw upon stored glycogen. We propose that G6P counteracts glycolysis in NP secreting neurons, thereby establishing a cellular environment that is more amenable to the expansion of the Golgi apparatus and large dense core vesicles, which in turn enhances NP release.

## RESULTS

### Identification of *G6P-GAL4* neuronal subtypes

Initial expression analysis of *G6P-GAL4; UAS-mCD8GFP* flies revealed the presence of approximately 14 GFP positive neurons in the brain (Miyamoto and Amrein, 2019). Three lines of evidence suggested that most, if not all, *G6P* expressing neurons were NP producing cells: First, they are relatively large in appearance and express DIMMED, a transcriptional regulator that plays a key role in cell fate determination of NP producing cells (Hewes et al., 2003). Second, one pair of cells located in the dorsomedial region co-expresses NPF, a *Drosophila* homolog of mammalian NPY. And third, examination of *G6P* expression in the thoracic ganglion showed many additional *G6P* positive cells, in a pattern similar to that described for neurons expressing FMRFamide and IPNamide (Lundquist and Nässel, 1990; Schneider et al., 1993b; Verleyen et al., 2004). These observations suggested that *G6P* is found in diverse populations of peptidergic neurons, expressing different NPs. We therefore set out to determine the identity of these NPs by comparing locations of G6P positive neurons with available reports of NP expression profiles, comprehensively reviewed by Nässel and Zandawala (Nässel and Zandawala, 2019). Based on this comparison, we carried out double staining experiments of brain and thoracic ganglia preparations of *G6P-GAL4*; *UAS-mCD8GFP* flies using NP specific antibodies and identified five NPs co-expressed with *G6P-GAL4* (Figure 1). In the brain, in addition to the prominent pair of *G6P^NPF^* neurons, Orcokinin-A, PDF and IPNa were found to be expressed in subsets of *G6P-GAL4* neurons (Figure 1A-1E, referred to as *Orco^G6P^*, *PDF^G6P^*, *IPNa^Br_G6P^* neurons), while in the thoracic ganglion, a group of FMRFa- positive neurons and a second subset of IPNa-positive neurons co-expressing *G6P-GAL4* were identified (Figure 1G-1I, referred to *FMRFa^G6P^* and *IPNa^Th_G6P^* neurons; note that the FMRFa- positive neurons present in the brain do not express *G6P-GAL4*; Figure 1F). Neurons expressing a given NP fall generally into two groups: a small number of neurons presenting with large cell bodies, and a larger number of neurons presenting with smaller cell bodies (Figure 1). For example, in the brain, there are some 20 NPF expressing neurons, only four of which fall into the large size class (Shao et al., 2017) (Figure 1J). Likewise, only two of the 10 Orcokinin positive neurons fall into the large class, while about half of the PDF positive neurons correspond to the large ventral lateral clock neurons (large LN-Vs) (Taghert et al., 2001). Of note, G6P is only expressed in large neurons, both in the brain and the thoracic ganglion (Figure 1B-1E, 1H and 1I). Together, about 30 of the 60 large neurons expressing the five NPs are G6P-positive (*NP^G6P^* neurons).

**Figure 1.**
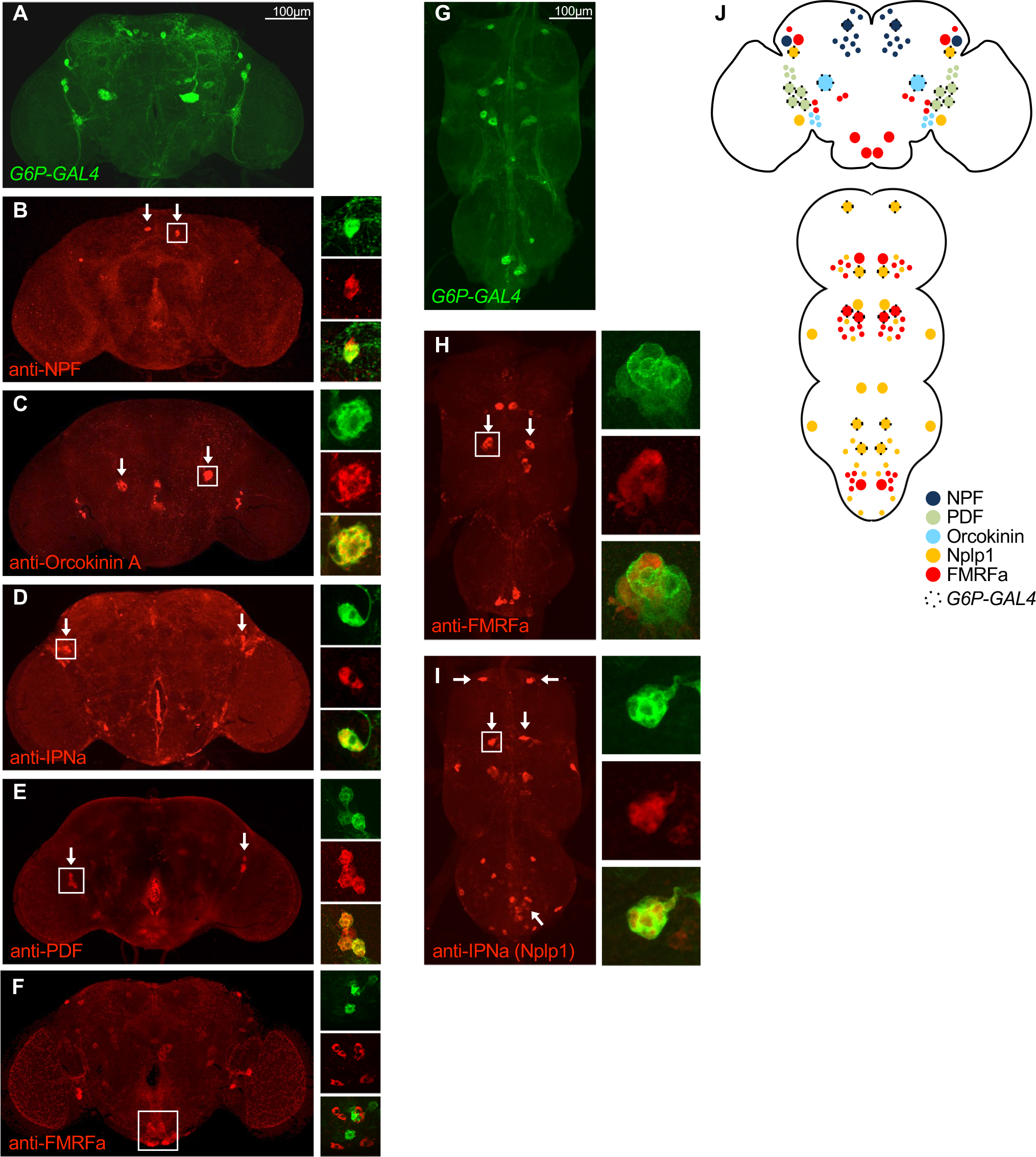
*G6P* is co-expressed in diverse subsets of peptidergic neurons. **A-I)** Antibody staining of the brain **(A-F)** and in the thoracic ganglia **(G-I)** of *G6P-GAL4; UAS- mCD8GFP* flies using anti-GFP antibody and antibodies against the indicated neuropeptides. *G6P-GAL4* is expressed in about 14 neurons in the brain, including a pair of large, centrally located NPF neurons **(B)** and a pair of large Orcokinin-A expressing neurons **(C)**, two IPNa **(D)** and four PDF neurons **(E)** which correspond I-LNvs. In the thoracic ganglia, *G6P-GAL4* and is expressed in about 10 to 16 neurons, including at least two pairs of FMRFa neurons **(H)** and three pairs of IPNa neurons **(I)**. **J)** Expression summary of *G6P-GAL4* neurons and relevant NP. Co-expressing neurons are outlined with a broken line. Note that all *G6P-GAL4* neurons are of the large variety.

In order to functionally manipulate subpopulations of specific NP expressing neurons, we procured *GAL4* lines expressed in Orcokinin, IPNa and FMRFa producing neurons from the *Flylight* repository (https://www.janelia.org/project-team/flylight) or obtained previously characterized lines for NPF (Wu et al., 2003) and PDF (Renn et al., 1999), respectively. We confirmed *GAL4* expression with these NPs by co-immunostaining brains and thoracic ganglia of *NP-GAL4; UAS-mCD8GFP* flies (Supplementary Figure 1).

### FMRFa peptides mediate glucose homeostasis

Expression of *G6P* in different NP secreting neurons suggests that G6P activity impacts several distinct NP signaling pathways. Using *G6P* mutant flies, we previously showed that one function of this enzyme is to maintain whole-body glucose homeostasis in starved flies, while having no significant impact on the levels of trehalose. Importantly, whole-body glucose homeostasis can be complemented in *G6P* mutant flies by temperature induced neural activation of *G6P* neurons (Miyamoto and Amrein, 2019) using *UAS-TRPA1* (Parisky et al., 2008). Similarly, silencing all *G6P-GAL4* expressing neurons using *UAS-Kir2.1* encoding the inward rectifier potassium channel Kir2.1 (Baines et al., 2001; Paradis et al., 2001) mimicked the whole-body glucose homeostasis phenotype of *G6P* mutant flies (Figure 2A)(Miyamoto and Amrein, 2019). To determine which subset of *NP^G6P^* neurons is responsible for glucose homeostasis, we expressed *UAS-Kir2.1* under the control of the five NP specific *GAL4* drivers. Inactivation of *NPF^G6P^*, *Orco^G6P^*, *PDF^G6P^*, *IPNa^Br_G6P^* or *IPNa^TG_G6P^* neurons caused no change in whole body glucose levels under either fed or starved conditions (Figure 2A). Only expression of *UAS-Kir2.1* in *FMRFa^G6P^* neurons (*GMR18C05-GAL4*) led to significantly reduced glucose levels in starved flies (Figure 2A). Consistent with our previous analysis, trehalose levels were not significantly different when compared to wild type control flies (Supplementary Figure 2). To test whether the function of *G6P* in *FMRFa* positive neurons is sufficient for maintaining whole body glucose homeostasis, we measured glucose levels in *G6P* mutant flies expressing a *UAS-G6P* transgene under the control of *GMR18C05-GAL4*. Indeed, these flies showed fully restored whole body glucose levels under starving conditions (Figure 2B). Taken together, these experiments identified *FMRFa^G6P^* neurons in the thoracic ganglion as the neurons responsible for maintaining whole body glucose homeostasis in food deprived flies.

**Figure 2.**
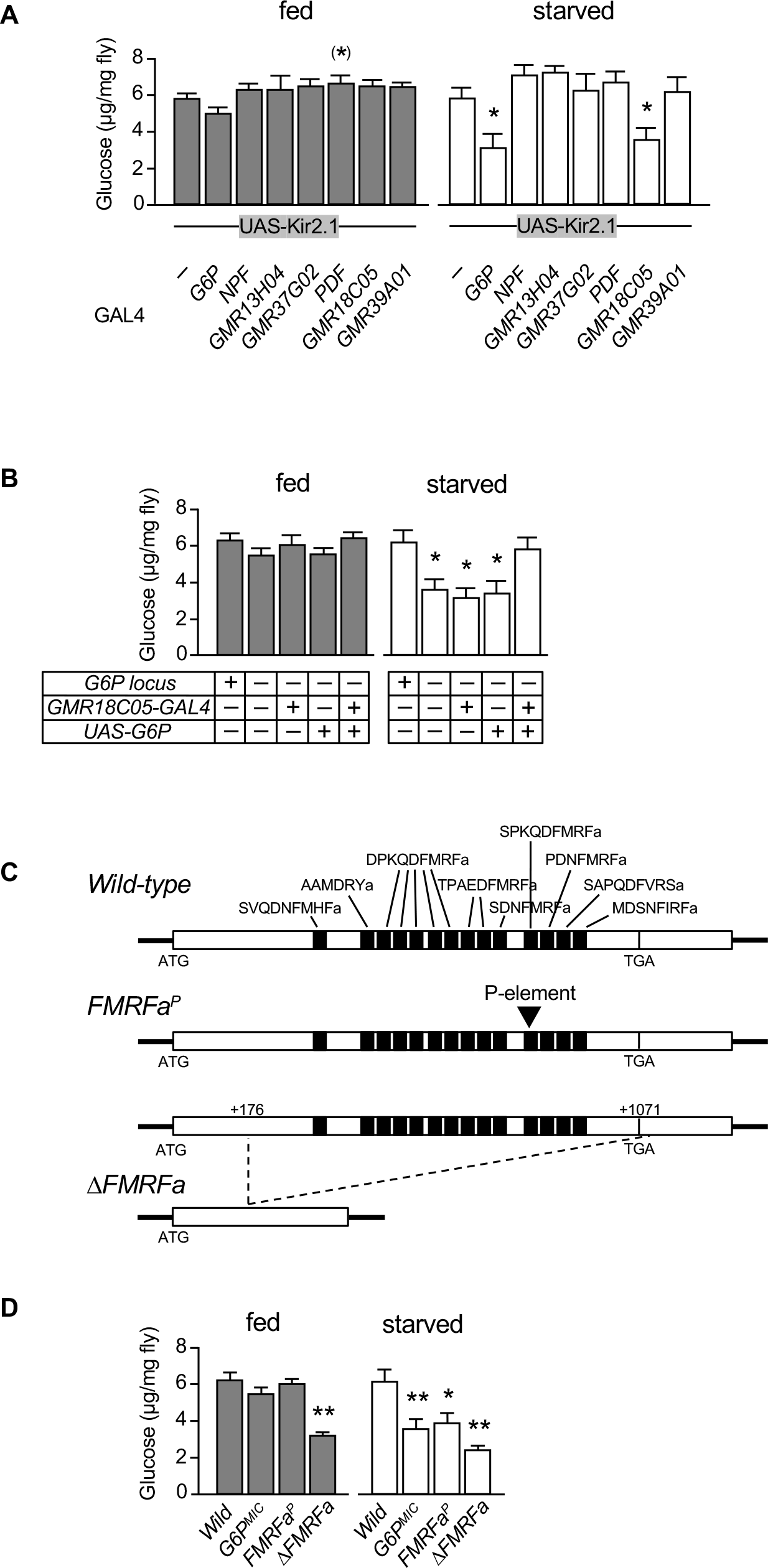
G6P mediates whole-body glucose homeostasis through FMRFa neurons. **A)** Silencing of all *G6P-GAL4* neurons or *GMR18C05-GAL4* (i.e. *FMRFa* expressing neurons; see Figure 3 and Supplementary Figure 1) using *UAS-Kir2.1*, but not silencing of any other subgroup, significantly reduces whole body glucose levels. *GMR13H04-GAL4* is expressed in *OrcoA^G6P^* neurons, *GMR37G02-GAL4* in *IPNa^Br_G6P^* neurons, *GMR39A01*-*GAL4* in *IPNa^Th_G6P^* neurons and *GMR18C05-GAL4* in *FMRFa^G6P^* neurons. Asterisks indicate *P<0.05 using Kruskal-Wallis with Dunn’s post-hoc test; n=8-10. Error bars represent standard error. **B)** *G6P* is required in *FMRFa* neurons for whole body glucose homeostasis in starved flies. Under fed conditions, both *wild-type* and *G6P* mutant flies maintain glucose levels (left). However, when starved for 24 hours, *G6P* mutant flies exhibit significantly reduced whole body glucose levels, a phenotype rescued when a *UAS-G6P* transgene is expressed in *FMRFa^G6P^* neurons under the control of *GMR18C05-GAL4*. Asterisks indicate *P<0.05 using Kruskal-Wallis with Dunn’s post-hoc test; n=9-12. Error bars represent standard error. **C)** Structure of the second, ORF containing exon of the wild type *FMRFa* gene and the *FMRFa[P]* and *ΔFMRFa* mutation. Exon 2 contains all 14 FMRFamides and FMRFamide-related peptides. The P-element in *FMRFa[P]* is inserted in the 11^th^ peptide, while all peptides are deleted in *ΔFMRFa*. The first, non-coding exon is not shown. **D)** FMRFamides are required for whole body glucose homeostasis. Like *G6P* mutants, *FMRFa[P]* mutants have significantly lower glucose levels in starved but not fed conditions. *ΔFMRFa* mutant flies show lower glucose levels regardless of feeding conditions. Asterisks indicate *P<0.05, **P<0.01 using one-way ANOVA with Kruskal-Wallis using Dunn’s post-hoc test; n=8-12. Error bars represent standard error.

*G6P-GAL4* exhibits varied expression in the thoracic ganglion. While consistently observed in three to four cells in the meso-thoracic segment, expression varied in the pro- and metathoracic segments. Thus, we sought to investigate overlap between *G6P* and *FMRFa* in more detail using the split GAL4 system, which reveals expression of a reporter only in cells that co-express two independent drivers controlled by the promotors of two genes of interest (Luan et al., 2006; Pfeiffer et al., 2010). We generated split-GAL4 transgenic flies (*G6P- GAL4DBD* and a *FMRFa-p65AD*) that also contained a *10xUAS-mCD8GFP* reporter and performed immunostaining of the CNS (Figure 3A and 3D). These experiments revealed that the thoracic ganglion harbored eight large, ventrally located *FMRFa^G6P^* neurons, which correspond to four pairs of Tv neurons (T1v, T2v, T2va and T3v; Figure 3A and 3C)(Schneider et al., 1993b). The same eight neurons are also labelled by *GMR18C05-GAL4* (Figure 3B and 3C), which we used for the functional characterization of *FMRFa^G6P^* neurons in all subsequent experiments (see below). In the brain, the split-GAL4 system and *GMR18C05-GAL4* did not label any FMRFa positive neurons (Figure 3D and 3E), confirming the observations obtained from double staining experiments using *G6P-GAL4* (Figure 1F).

**Figure 3.**
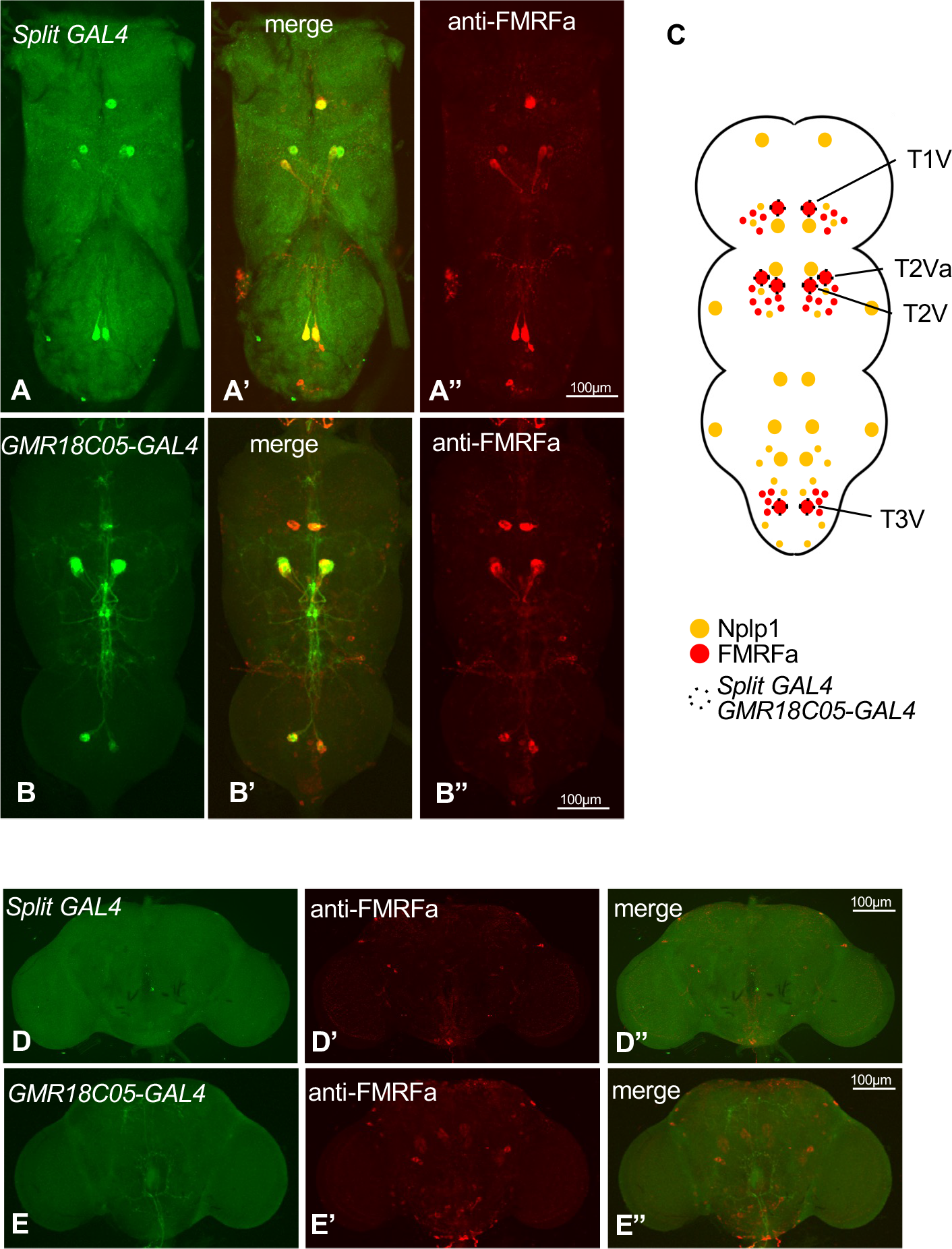
***FMRFa^G6P^* neurons correspond to 8 Tv neurons in the thoracic ganglion**. The split GAL4 system (*G6P-GAL4DBD* and *FMRFa-p65AD*) was used to identify neurons that express both *G6P* and *FMRFa.* The CNS of *G6P-GAL4DBD/FMRFa-p65AD; 10xUAS-mCD8GFP* flies **(A and D)** and *GMR18C05-GAL4; UAS-mCD8GFP* flies **(B and E)** was dissected and stained with anti-GFP and anti-FMRFa antibody. **A-B)** Eight neurons in the thoracic ganglion were labeled by the split GAL4 drivers, all of which also express FMRFa (A’ and A”). These neurons are also labeled by *GMR18C05-GAL4* (B, B’ and B”). **C)** Expression summary of GAL4 drivers in the thoracic ganglion. The eight large FMRFa neurons labeled by either the split GAL4 system or *GMR18C05-GAL4* correspond to the Tv neurons shown to express FMRFa (Schneider et al., 1993b). **D- E)** Neither split GAL4 nor *GMR18C05-GAL4* are expressed in any FMRFa neurons in the brain.

*FMRFa* encodes a 347 amino acid long propeptide, which is processed into 14 peptides, 10 of which are characterized by the carboxyterminal FMRF motif (Figure 2C) (Predel et al., 2004). To further confirm that FMRFamides are the signaling molecules mediating glucose homeostasis in starved flies, we measured glucose levels in *FMRFa^P^* mutant flies. *FMRFa^P^* is a P-element insertion into the 11^th^ of the 14 peptides and homozygous *FMRFa^P^* mutant, starved flies showed the same glucose homeostasis deficit as flies in which these neurons were silenced (Figure 2D). We also generated a CRISPR/Cas-9 null allele, a deletion encompassing all 14 peptides (*ΔFMRFa*, Figure 2C, Supplementary Figure 3), and found that *ΔFMRFa* flies showed reduced whole body glucose levels in both fed and starved conditions (Figure 2D).

### G6P is necessary to maintain glucose levels in *FMRFa^G6P^* neurons

*NPF^G6P^* neurons in the brain require G6P cell-autonomously to maintain intracellular glucose levels when flies are starved (Miyamoto and Amrein, 2019). In these experiments, we employed the fluorescent sensitive glucose sensor Glu700 and found that *NPF^G6P^* neurons of starved *G6P* mutant flies were unable to maintain intracellular glucose levels (Miyamoto and Amrein, 2019). While the relevance of this phenomenon for proper neuropeptide signaling remains to be determined, the glucose sensor provides a means to assess whether G6P is also necessary for maintaining intracellular glucose levels in *FMRFa^G6P^* neurons during starvation. Thus, we established an *in vivo* preparation of the ventral nerve cord that allowed us to monitor glucose levels in these neurons (see Material and Methods). Since G6P is an Endoplasmic Reticulum (ER) resident enzyme (van Schaftingen and Gerin, 2002), we expressed an ER resident version of *Glu700, Glu700KDEL* (Addgene, #17867) under the control of *GMR18C05-GAL4*. In fed *G6P* mutant flies, glucose levels were indistinguishable from wild type fed flies, while in starved flies, *G6P* mutants showed significantly lower glucose levels in *FMRFa^G6P^* neurons compared to fed flies (Figure 4). Together, these results show that in starved flies, G6P is also functional in *FMRFa^G6P^* neurons where it maintains intracellular glucose levels during starvation.

**Figure 4.**
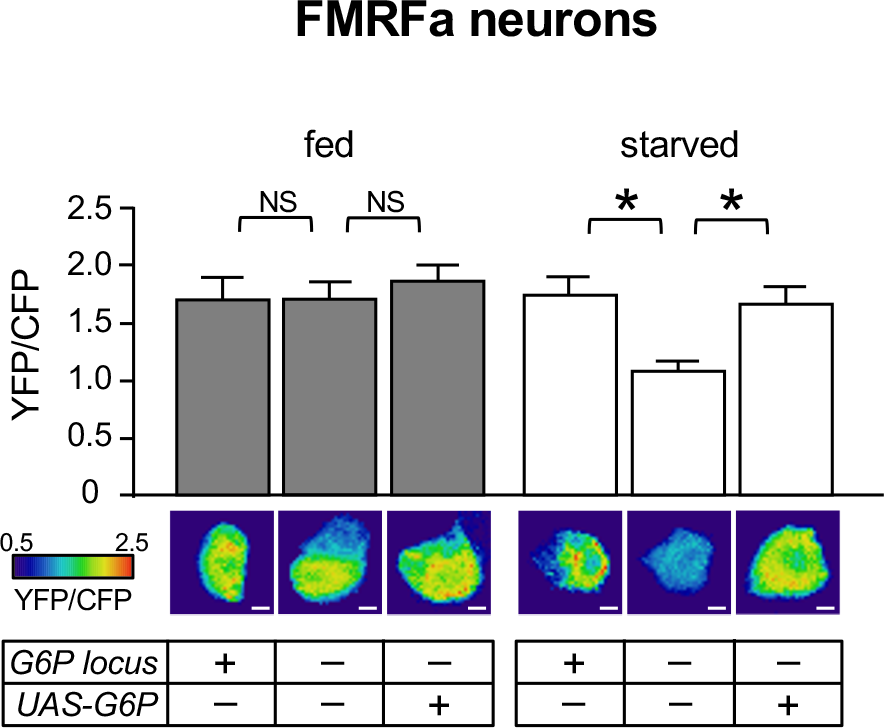
***G6P* is required to maintain glucose levels in FMRFa neurons of food-deprived flies**. Glucose levels were measured using Glu700KDEL, a FRET-based, ER tagged glucose sensor, in FMRFa neurons in the thoracic ganglion of *GMR18C05-GAL4/UAS-Glu700KDEL* flies. Glucose levels are reduced in starved *G6P* mutant flies, which was rescued in the presence of a *UAS- G6P* transgene. Asterisk indicates **P<0.01 using one-way ANOVA with Tukey post hoc test. N=8-17. Error bars represent standard error.

### G6P increases Golgi network volume and facilitates NP release

NP processing involves propeptide cleavage and transport through the Golgi network, ultimately leading to NP deposition in Large Dense Core Vesicles (LDCVs) that derive from the trans Golgi stack (van den Pol, 2012). Thus, we examined whether any difference in size and overall distribution of the ER and Golgi network of *FMRFa^G6P^* neurons was apparent between *wild type* and *G6P* mutant flies using ER (GFP-KDEL)(Kneen et al., 1998) and Golgi (αManll- GFP) resident reporter proteins (Hardt et al., 2005; Zhou et al., 2014). No obvious differences were observed in the ER. However, the Golgi network occupied a significantly smaller area in *G6P* mutant flies compared to *wild type* flies (Figure 5A 9% vs 14%; see also Supplementary Figure 4). A similar reduction in Golgi volume was also observed when these reporters were expressed in the pair of *NPF^G6P^*neurons (9% vs 16%, Figure 5B; left panels). Of note, and in contrast to mammalian cells where the Golgi complex is organized as a single ribbon, the *Drosophila* Golgi apparatus is dispersed throughout the cytoplasm (Kondylis and Rabouille, 2009).

**Figure 5.**
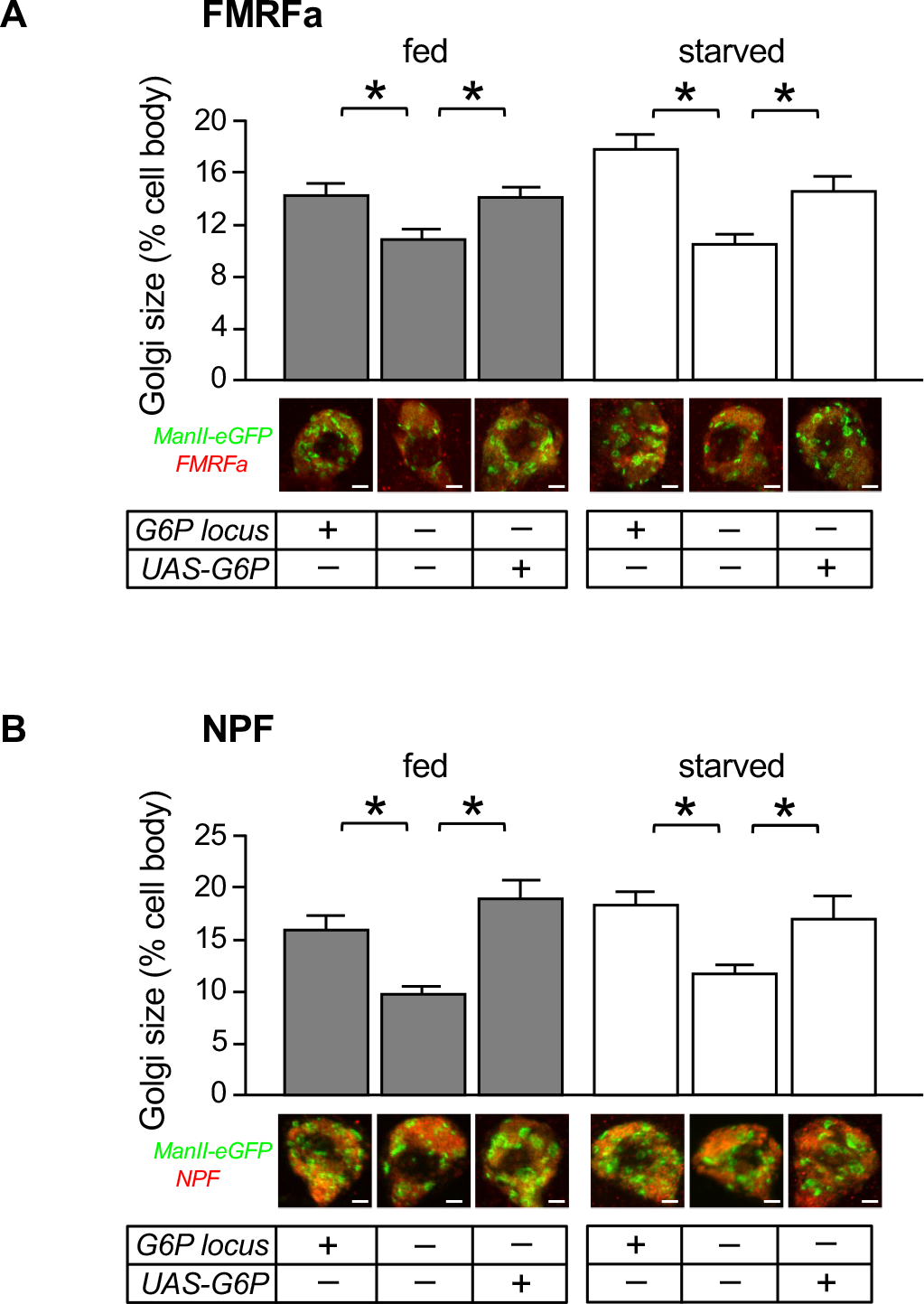
G6P positively affects the size of Golgi apparatus in the peptidergic neurons. The Golgi apparatus was visualized by expressing the GFP tagged Golgi marker UAS-ManII- eGFP. FMRFa staining was used as a proxy for cell body. Golgi size is given as percentage of cell body size (see also Supplementary Figure 4). *FMRFa^G6P^* neurons were identified using *GMR18C05-GAL4* driving *UAS-ManII-eGFP* expression **(A)** and *NPF^G6P^* neurons were identified using *NPF-GAL4* driving *UAS-ManII-eGFP* expression **(B)**. Neurons of *G6P^-/-^* flies have a significantly smaller Golgi apparatus, in fed and starved conditions. This phenotype is rescued *UAS-G6P* transgene expression. Asterisks indicate *P<0.05 using Kruskal-Wallis with Dunn’s post-hoc test. N=44-65 for *FMRFa^G6P^* neurons, N=12-23 for *NPF^G6P^* neurons. Error bars represent standard error. Scale bars are 4µm.

Because LDCVs emerge from the trans Golgi network, NP secretion might be negatively affected in *G6P* mutant flies. To measure the effect of G6P on NP release, we took advantage of the fact that the *FMFRa^G6P^* neurons correspond to the Tv neurons. These neurons innervate the neurohemal like area in the dorsal neural sheet of the thoracic ganglion, which are release sites into the hemolymph (Lundquist and Nässel, 1990; Schneider et al., 1993b). Thus, NP release from *FMRFa^G6P^* neurons can be assessed by measuring the NP amounts in the hemolymph. Because available antibodies against FMRFa cross-react with other hemolymph components/peptides, we used the mammalian Atrial natriuretic peptide (Anf) fused to GFP as a proxy NP (Anf-GFP, Husain and Ewer, 2004) and assessed Anf-GFP amounts released into the hemolymph from *FMRFa^G6P^* neurons of *wild-type* and *G6P* mutant flies, as well as the total amount of Anf-GFP in these flies (Figure 6A and 6B). It was previously established that Anf-GFP is released in a time-appropriate fashion during molting, following the dynamic profile of the resident peptides in these cell, Ecdysis Triggering Hormone (Husain and Ewer, 2004). We expressed the *UAS-Anf-GFP* reporter gene in *FMFRa^G6P^* neurons and measured peptide levels in the hemolymph. Compared to the *wild-type*, *G6P* mutant flies secrete significantly lower amounts of Anf-GFP into the hemolymph in both fed and starved flies (Figure 6A). Importantly, the deficit can be rescued by inclusion of a *UAS-G6P* transgene. Total amount of Anf-GFP was not affected by *G6P* or lack thereof, but starved flies contained about 30% more NP compared to fed flies (Figure 6B). To explore whether *G6P* dependence can be imposed on non-*G6P* neurosecretory cells, we measured hemolymph and total levels of an epitope tagged, Insulin like peptide 2 (Ilp2), expressed from a genetically modified *Ilp2* gene (*Ilp2^1^ gd2HF*) (Park et al., 2014). When *UAS-G6P* was expressed in insulin producing cells (IPCs; *Ilp2^1^ gd2HF/dilp2-GAL4 UAS-G6P*), a significant increase of secreted Ilp2HF was observed in starved flies, with a concomitant decrease in total Ilp2HF (Figure 6C and 6D; white bars). Given that IPCs of starved flies do not actively secrete Ilp2HF (Park et al., 2014), this observation implies that ectopic expression of *G6P* triggers quiescent IPCs to secrete Ilp2HF. Total amount of Ilp2HF in fed flies was slightly increased in the presence of G6P, which however, did not result in an increase in secreted Ilp2HF (Figure 6C and 6D; black bars). Furthermore, fed and starved control flies (i.e. flies not expressing G6P) exhibited similar levels of hemolymph Ilp2HF (Figure 6C), an observation that appears counterintuitive based on observations made by Park and colleagues, who reported that hemolymph Ilp2HF spikes rapidly after a meal given to previously starved flies. However, this spike is transient and recedes to levels of starved flies within 60 minutes. (Park et al., 2014). In our experimental set up, fed flies have constant access to food, allowing for a wide range of different feeding states within the cohort at the time of hemolymph collection. Thus, similar amounts of hemolymph Ilp2HF in fed and starved control flies is probably due to the fact only a small fraction of flies, if any, have taken a meal prior to hemolymph collection. Regardless, the observations made in starved flies argues that ectopic expression of *G6P* in IPCs enhances their capabilities to secrete Ilp2HF.

**Figure 6:**
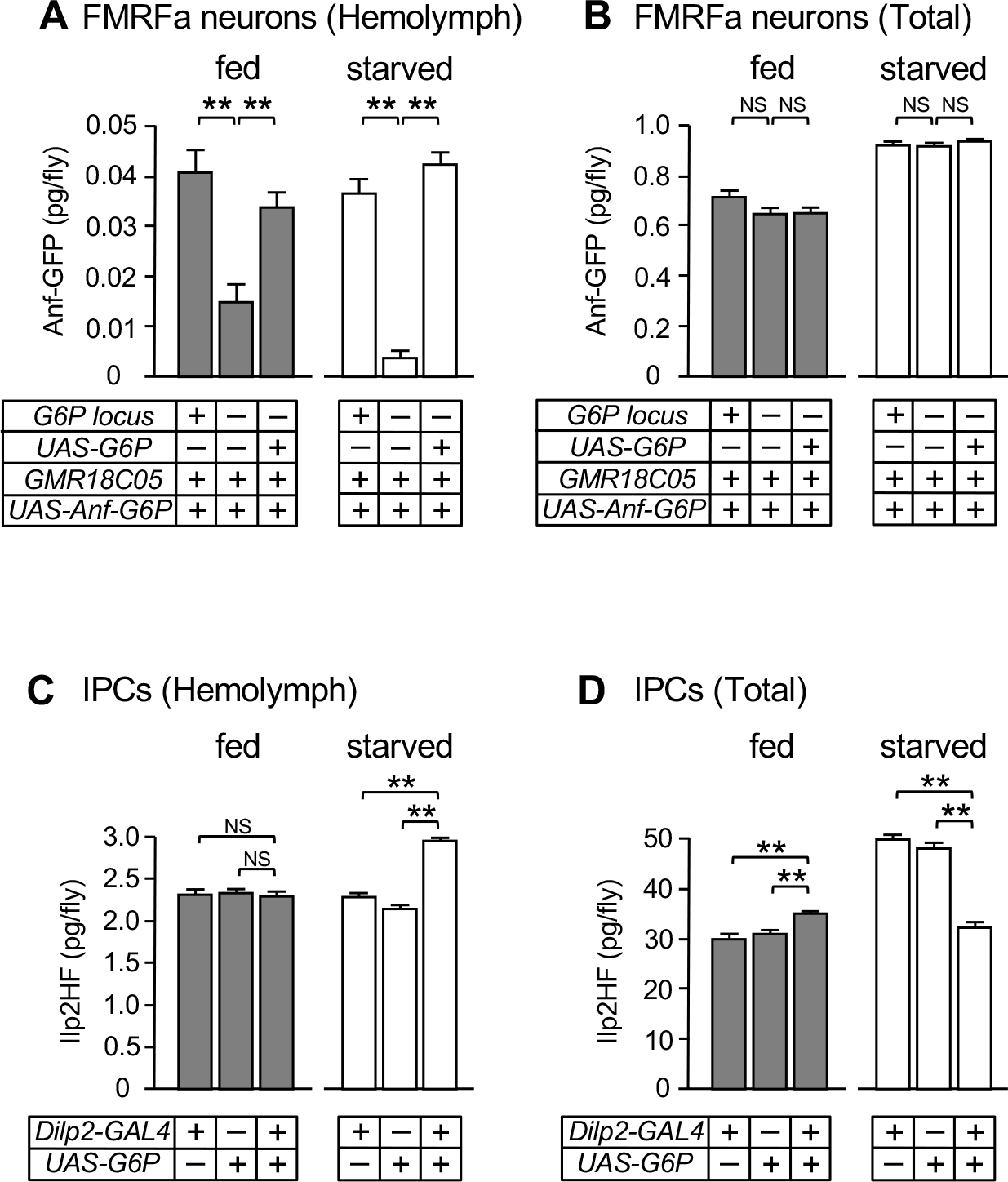
FMRFa neurons require G6P for efficient neuropeptide secretion. **A-B)** Secreted **(A)** and total amount **(B)** of Anf-GFP was quantified by ELISA (Rat ANP ELISA Kit; abcam, ab108797) in flies expressing *UAS-Anf-GFP* in *FMRFa* neurons (*GMR18C05-GAL4*). Compared to *wild-type* (left bar in each set), the hemolymph of *G6P* mutant flies (center bar) contains significantly lower amounts of Anf-GFP in both the fed and starved flies **(A)**. Secretion of Anf-GFP is rescued in the presence of a *UAS-G6P* transgene (right bar in each set. Total amount of Anf-GFP was unaffected by the presence or absence of G6P **(B)**. **C - D)** Secreted **(C)** and total amount **(D)** of Ilp2HF was quantified in *Ilp2^1^ gd2HF* flies, which carry a genetically tagged Ilp2 allele. Ectopic expression of *UAS-G6P* under the control of *dilp2-GAL4* increased Ilp2HF levels in the hemolymph of starved flies **(C)** with concomitant reduction in total Ilp2HF **(D)**. In fed flies, a small increase of total Ilp2HF was observed. Asterisks indicate **P<0.01 using Kruskal-Wallis with Dunn’s post-hoc test. N=12 for *FMRFa^G6P^* neurons. N=9 for IPC. Bars represent standard error.

### *G6P* mediated FMRFa signaling to the jump muscle for glycogen storage

In invertebrates, FMRFamides have been associated with the modulation of heart rate, gut motility, sleep, flight and reproduction (Brussaard et al., 1988; Fisher et al., 1993; Lenz et al., 2015; Nässel and Zandawala, 2019; Ravi et al., 2018). Yet, the roles of FMRFamides and their receptors remains incomplete, in part because many of these studies have been conducted in systems not amenable to molecular genetic analyses. Thus, we sought to determine the target tissue of FMRFamides, expected to express its receptor (FMRFaR, CG2114), and to identify phenotype(s) in flies carrying *FMRFaR* mutations. We took advantage of the chemoconnectomics resource generated by Deng and colleagues, who engineered a series of *GAL4* knock-in lines for many NP receptor genes, including *FMRFaR* (Deng et al., 2019). In adult flies, *FMRFaR^2A-GAL4^* is strongly expressed in the jump muscle and the CNS, while no expression was found in other tissues (flight muscle, reproductive system, fat body, gut etc; Figure 7A), a finding consistent with expression profiles reported in the aging fly cell atlas (Lu et al., 2023). We acquired a *FMRFaR* mutant strain (*FMRFaR^MB04659^*) and examined the amount of muscle glucose and glycogen in *wild type* and homozygous receptor mutant flies (Figure 7B). While free glucose levels were not noticeably different in the jump muscle, regardless of feeding status, muscle glycogen was significantly lower in fed *FMRFaR^MB04659^* homozygous mutant flies, when compared to fed wild type flies (Figure 7B). *FMRFaR^MB04659^* mutant flies, regardless of feeding status, exhibited similar glycogen levels as starved wild type flies, which amounts to about 50 percent of that observed in fed wild type flies. Glycogen loss was fully rescued in the presence of a *UAS-FMRFaR* transgene driven by *Act79B-GAL4* (Figure 7C), which is predominantly expressed in the jump muscle (Bryantsev et al., 2012). Importantly, the presence of *ELAV-GAL80* in these flies eliminates the potentially confounding contribution of low level *Act79B-GAL4* expression in the CNS. Moreover, ectopic expression of *FMRFaR* in the fat body, in contrast to expression in the jump muscle, does not restore glycogen levels in *FMRFaR* mutant flies (Figure 7C). And lastly, we note that glycogen levels were also significantly reduced in fed *G6P* and *FMRFa* mutant flies (Figure 7D). Taken together, these results indicate that one key function of FMRFa signaling is the generation of glycogen stores in the jump muscle, which can be accessed to generate glucose in times of limited nutrient availability.

**Figure 7:**
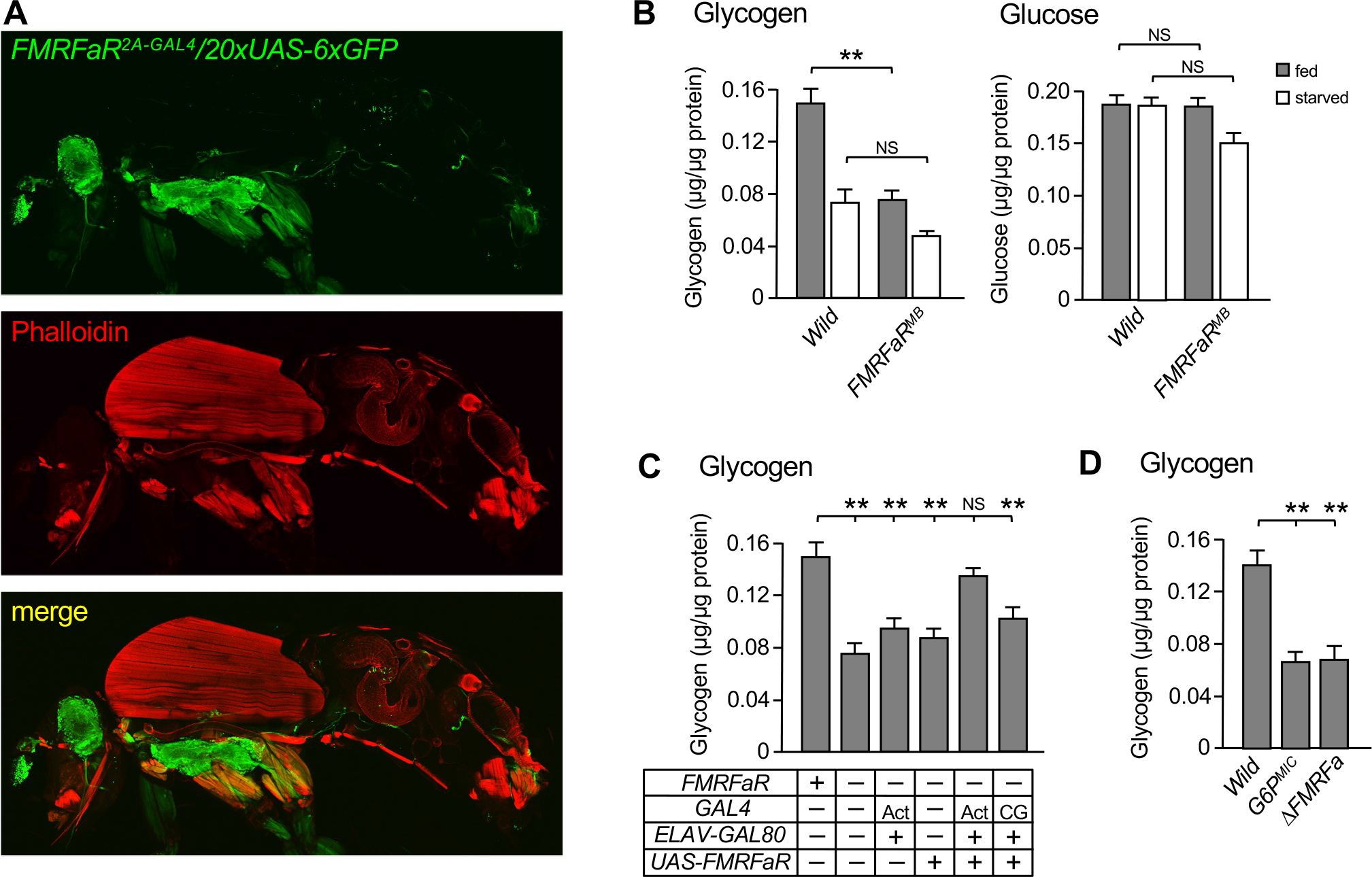
FMRFa acts on the FMRFa receptor to establish glycogen stores in the jump muscle **A)** *FMRFaR* was visualized using a *GAL4* gene knock in allele (*FMRFaR^2A-GAL4^*) combined with *UAS-20xUAS-6xGFP*, following anti-GFP staining. Flies show robust expression in the jump muscle and CNS. Preparation was counterstained with Phalloidin (Invitrogen A34055). **B)** Glycogen stores in the jump muscle were significantly smaller in the *FMRFaR^MB^* mutant flies under starved conditions, while glucose levels remained unchanged. **C)** Glycogen stores were restored to *wild type* levels when *UAS-FMRFaR* was expressed in the jump muscle in *FMRFaR^MB^* mutant flies also containing *ELAV-GAL80* suppressor. Expression of *UAS-FMRFaR* in the fat body using the *CG-GAL4* driver, however, did not restore glycogen levels in the jump muscle. **D)** Glycogen loss in the jump muscle is also observed in the *G6P^MIC^* and *ΔFMRFa* mutant flies. Asterisks indicate *P<0.05, **P<0.01 by one-way Kruskal-Wallis with Dunn’s post-hoc test. n=7-14. Error bars represent standard error.

## DISCUSSION

Activity of NPs depends on many external and internal factors, providing a means for temporally and quantitatively controlled release that accommodates an animal’s physiological demands. In addition to regulation of transcription and RNA processing (Gauthier and Hewes, 2006; Herzog et al., 1997; Higuchi et al., 2005; Schneider et al., 1993a), most NPs undergo post-translational modification through proteolytic cleavage from precursor proteins (pro-peptides)(Hook et al., 2008; Kovac et al., 2009; Pauls et al., 2014; Rholam and Fahy, 2009). Generally, these regulatory processes are specific to NPs, as they must be tailored to sequence and/or structural features within the specific NP gene or protein. The studies presented here report on a new mode of regulation in NP signaling, namely one that allows modulation across distinct types of secretory neurons expressing the gluconeogenic enzyme G6P. Based on measurements of hemolymph Anf-GFP peptide released from *FMRFa^G6P^*neurons and Ilp2HF secreted by IPCs ectopically expressing a *UAS- G6P* transgene, we postulate that G6P primes peptidergic neurons for more efficient NP release, regardless of the NP they express. Thus, it will be interesting to explore whether G6P modulates NP release in other *NP^G6P^* neurons and affects respective physiological processes and behaviors, such as courtship (NPF; Liu et al., 2019; Silva et al., 2021), oogenesis (Orcokinin, Silva et al., 2021), circadian behavior and sleep (PDF, Chung et al., 2017; Lear et al., 2009; Parisky et al., 2008), or feeding (NPF, Landayan et al., 2021; Lear et al., 2009; Lee et al., 2004). To our knowledge, a function for an enzyme with an established role in gluconeogenesis and glycogenolysis in neurosecretion, or neurons in general, is unprecedented. While a specific mechanism by which G6P modulates peptidergic neurons remains to be determined, the diminished Golgi size in mutant flies implies that G6P activity creates a cellular environment that is beneficial to Golgi expansion (see below).

### G6P increases Golgi apparatus and enhances NP release

The central function of mammalian G6P is to catalyze hydrolysis of g-6-p during glycogenolysis or gluconeogenesis, generating free glucose that is exported and released into the blood for systemic glucose homeostasis when animals are food deprived. Despite the fact that *Drosophila* G6P has such catalytic activity in peptidergic neurons and is required to maintain systemic whole-body glucose levels (Miyamoto and Amrein, 2019), multiple lines of evidence argue against a role for *Drosophila* G6P to produce glucose for systemic sugar homeostasis. First, we have previously shown that glucose homeostasis can be restored in starved *G6P* mutant flies by expression of TRPA1 in *G6P* neurons, arguing for a signaling role (Miyamoto and Amrein, 2019). Second, neurons in general have high energy demands and are ill equipped for the generation of glucose, for which major amounts of stored glycogen or incidental supply of amino acids is required. Third and related to this point, while *G6P* peptidergic neurons are diverse, their primary purpose is to produce and secrete NPs, an energy costly process that seems difficult to reconcile with other energy consuming activities such glycogenolysis or gluconeogenesis, although fat cells, which carry out both glycogenolysis and trehaloneogenesis are also prolific hormone producers. And lastly, in nutrient deprived flies, the fat body is well equipped to maintain sugar homeostasis by generating trehalose from g-6-p. These observations, and the fact that neuronal glycolysis is largely dispensable (see below), lead us to posit that the function of *G6P* is to counteract glycolysis in peptidergic neurons, creating conditions that are overall beneficial to the main task of these neurons, the release of NPs.

Volkenhoff and colleagues showed that glycolysis is dispensable in neurons (Volkenhoff et al., 2015). Specifically, pan-neuronal RNAi knock down of numerous glycolytic enzymes in the CNS had no impact on survival and overall locomotion, while ubiquitous or pan-glial knock-down of the same genes led to lethality or severe locomotion deficits. Thus, glial but not neuronal glycolysis is essential in the CNS for survival. Moreover, this study also established that lactate generated in glia is shuttled to neurons, where it is used for ATP production via the TCA cycle. While neurons are likely equipped with glycolytic enzymes necessary for energy production, it is apparent that at least in *Drosophila*, neurons rely mostly on metabolites from glia that drive alternative metabolic processes. Based on these findings and the observations reported in this paper, we posit that inhibition of glycolysis in *NP^G6P^* neurons creates a cellular environment that enhances production, transport and/or secretion of NPs. For example, shutdown of glycolysis reduces the amounts of glycolytic metabolites, which will affect the intracellular pH and hence might increase efficacy of numerous biochemical processes beneficial for the expansion of the Golgi apparatus and ultimately for NP transport and/or secretion.

### Diverse roles for G6P enzymes

Humans and most mammals have two additional genes encoding G6P, *G6PC2* expressed in the pancreas, Müller glia cells and spermatids, and the ubiquitously expressed *G6PC3*. *G6PC2* deficient mice show lower fasting blood glucose (Bosma et al., 2020), while humans with mutations in *G6PC3* present with heart defects and neutropenia (Banka et al., 2011; Cheung et al., 2007). An intriguing underlying cellular mechanisms for neutropenia in patients with mutations in *G6PC3* has been proposed (Veiga-da-Cunha et al., 2019), namely the accumulation of 1,5-anhydroglucitol-6-phosphate (1,5AG6P), a toxic metabolite derived from a common circulating food compound (1,5-anhydroglucitol). This is consistent with the observation that *G6PC3^-/-^* neutrophils express numerous markers indicative of ER stress, while also exhibiting glycosylation defects (Hayee et al., 2011). Thus, it appears that the main role of G6PC3 in neutrophils is to remove a metabolite that is detrimental to optimal physiological processes within these cells. Alternative - and potentially neuronal - roles for G6P enzymes are also indicated by evolutionary arguments, because the *G6P* genes came into existence in Cnidaria, one of the first animal lineages that possessed neuronal cell types, but lacked organs comparable to the liver or even a basic circulatory system (Miyamoto and Amrein, 2017).

### FMRFamide signaling is essential to build up muscle glycogen stores

Our investigations have identified a primary function for FMRFamides in glycogen metabolism. Under ad libitum feeding conditions, *wild type* flies establish considerable glycogen stores, which can be mobilized in the form of glucose when food is scarce. In contrast, *G6P^MIC^*, *ΔFMRFa* and *FMRFaR^MB04659^* mutant flies generate only about half the amount of glycogen in the jump muscle (Figure 7B) and consequently are unable to maintain whole body glucose homeostasis when deprived of food. Thus, FMRFa signaling is essential during normal feeding conditions to assure that glycogen stores are established in the jump muscle.

Sugar homeostasis is remarkably well-conserved between mammals and insects. While in mammals, where counteracting hormones - insulin and glucagon – are secreted from pancreatic β and α cells to maintain blood sugar balance, fruit flies use counterbalancing peptides secreted from peptidergic neurons in the CNS for the same process - ILPs and adipokinetic hormone (AKH, the fly ortholog of mammalian glucagon)(reviewed by Kim et al., 2021). An increase in circulating sugars or lipids triggers insulin and ILP secretion from β cells and the IPCs, respectively, which in turn leads to the build-up of fat and glycogen stores in muscle and adipose tissue. When blood and hemolymph sugar levels are low, glucagon and AKH secretion from pancreatic α cells and the Corpora Cardiaca (CC) increases, which ultimately leads to the mobilization of stored fat and glycogen in mammals and insects, respectively. In the fly, the secretion of ILPs and AKH is coordinated by the CN neurons that directly communicate with IPCs and CC neurons, activating the former while suppressing the latter in response to high brain hemolymph glucose levels (Oh et al., 2019). While no direct neural connections between *FMRFa^G6P^* neurons and IPCs, CC or CN neurons are apparent, it has become evident that multiple, possibly independent, NP signaling pathways control carbohydrate metabolism. Indeed, a third brain sugar sensor was identified in the posterior superior lateral protocerebrum (Miyamoto et al., 2012). This sensor is activated by a spike of the low abundant hemolymph sugar fructose, via the evolutionarily conserved gustatory receptor Gr43a (Fujii et al., 2023). These neurons, which intriguingly express yet another NP (corazonin; Miyamoto and Amrein, 2014), trigger opposite feeding behaviors, promoting food intake when flies are starved, but suppressing food intake when satiated.

Our analysis has shown that *FMRFaR* expression is restricted to the jump muscle and the CNS, an observation supported by the “aging fly atlas” (Lu et al., 2023), but this does not exclude the possibility that other tissues might be modulated by FMRFamides. For example, indirect signaling from the jump muscle might lead to larger glycogen stores in other tissues such the indirect flight muscle or the adipose, or FMRFamides might signal to yet unknow peptide receptors. In this context, we note that both *G6P* and *FMRFa* mutant flies, but not *FRMFaR* mutant flies, exhibit impaired climbing ability, suggesting that the *FMRFa^G6P^* neurons might impact metabolic processes through *FMRFaR* independent signaling pathways (Supplementary Figure 5). Indeed, Song and colleagues showed that a cysteine rich protein diet upregulates FMRFa signaling, which promotes energy expenditure and reduces food intake, resulting in overall loss of fat in the adipose tissue (Song et al., 2023). Their paradigm was based on dietary intervention (high cysteine diet), while our study investigated the function (and loss thereof) of *FMRFa* signaling under normal feeding and starvation conditions. It will be interesting to see whether, and if so, how different metabolic roles of *FMRFa* are interconnected. In another recent report, Ravi and colleagues identified a role for the FMRFaR in the CNS, establishing its requirement in dopaminergic neurons for sustained flight. However, possible metabolic deficits in these flies or the involvement of FMRFamides as signaling molecules was not investigated. Yet, it is likely that energy for sustained flight also requires stored glycogen, provided from the indirect flight muscle or other organs, such as the adipose. Indeed, muscle has gained considerable attention as a metabolic hub communicating with other organs, including the brain, gut and adipose tissue (Demontis et al., 2013, 2013; Gomarasca et al., 2020; Rai and Demontis, 2016; Zhao et al., 2020; Zhao and Karpac, 2017). Thus, our study and those by Ravi and Song and colleagues have implicated *FMRFa* signaling as one likely source in the regulation of different, yet possibly intersecting, nutrient metabolic signaling processes across different organ systems.

## ACKNOWLEGEMENTS

We thank Dr. Paul Hardin for the *Pdf-GAL4* strain, Dr. Gaiti Hasan for the *UAS-FMRFaR* strain, Dr. Richard Cripps for the *Act79B-GAL4* strain and Dr. Seung Kim for *Ilp2^1^ gd2HF* strain. We also thank Dr. Wolf Frommer for the Glu700KDEL plasmid (Addgene plasmid #17867) and Dr. Simon Bullock for pCFD3 plasmid (Addgene plasmid #49410), Dr. Gerald Rubin for the GAL4DBD and p65AD plasmids (Addgene #26233 and #26234). Antibody gifts were provided by Dr. Mizoguchi (anti-Orcokinin), Dr. Thor (anti-IPNa, anti-proFMRFa) and Dr. Hardin (anti-PDF). We thank Anastasia Lucas and Edward Wang for assistance with glycogen measurements and Drs. Dick Nässel and Raquel Sitcheran for valuable comments on the manuscript. We are grateful to Anastasia Lucas and Edward Wang for assistance on the metabolic assays. This work was supported by an NIH grant 1R21NS118118-01 awarded to H.A.

## MATERIAL AND METHODS

### Fly strains and maintenance

Experiments were performed using adult *Drosophila melanogaster* males, aged between 7- 14 days. Flies were reared on the standard fly food containing cornmeal, yeast and sugar at 23-24°C. *G6P^MI12250^* is referred as *G6P^MIC^*, *FMRFa^KG01300^* is referred as *FMRFa^P^*, *FMRFaR^MB04659^* is referred as *FMRFaR^MB^*. See supplementary data for detailed genotypes.

### Quantification and Statistical Analysis

Statistical significance was calculated using Student’s t-test or two-tailed Mann-Whitney U test for pairwise comparisons, and one-way ANOVA with Tukey post hoc test or Kruskal-Wallis test with Dunn’s post-hoc test for multiple comparisons. Error bars are standard error of the mean (SEM) except where noted. Significance: *p<0.05, **p<0.01, NS: not significant. All replicates are from different biological samples (testing different flies or cells). Image analysis was performed with NIS-Elements software suite. Statistical analysis was performed in Real Statistics Add-Ins in MS Excel and GraphPad PRISM. Number of replicates was 8-10 (Figure 2A), 9-12 (Figure 2B), 8-12 (Figure 2D), 12-17 (Figure 4), 51-65 (Figure 5A), 12-23 (Figure 5B), 11 (Figure 6A), 9 (Figure 6B), 7-14 (Figure 7B), 12-14 (Figure 7C), 7 (Figure 7D).

### Molecular Biology

The *UAS-FLII^12^Pglu-700µ*δ*6KDEL* was generated from the expression plasmid pEF/myc/ER *FLII^12^Pglu-700µδ6* (Addgene). The glucose sensor coding sequence was excised using PstI and XbaI. This fragment, along with a fragment encoding an ER targeting module (5’- GAACA<u>GAATTC</u>AGATCTGCCACC**ATG**GGATGGAGCTGTATCATCCTCTTCTTGGTAGCAACAGCTAC AGGCGCGCACTCCCAGGTCCAA<u>CTGCAG</u>TCA-3’) cut with <u>EcoRI</u> and <u>PstI</u>, were cloned in a three way cloning step into the *pUAST* vector, cut with EcoRI and XbaI. This construct was injected into *w^1118^* embryos (Bestgene, Inc). For construction of *G6P-GAL4DBD*, the promoter region of *G6P* was amplified from genomic DNA of the *w^1118^* strain, using 5’- GGCGGCCGCCACTTCGAGAACTTTGGATAGGT-3’ and 5’-GGGTACCATCGACTCCAATGTTTTCAGTCTT-3’ primers. DNA fragment was cloned into *pBPZpGAL4DBDUw* vector (addgene #26233)(Pfeiffer et al., 2010).

For construction of the *FMRFa-p65AD*, the promoter region of *FMRFa* was amplified using 5’- CGTTAACGTTCGAGGTCGACTCTGCAGACGTGGTTTTCG-3’ and 5’- CGTTAACGTTCGAGGTCGACTCTGCAGACGTGGTTTTCG-3’ primers (Schneider et al., 1993a).

DNA fragment was cloned into pBPp65ADZpUw (addgene #26234) (Pfeiffer Rubin 2010). All constructs were injected into *w^1118^* embryos (Bestgene, Inc).

Generation of *FMRFa* null mutations: *FMRFa* mutation was generated using CRISPR/Cas9 mutagenesis. The guide RNA (gRNA) was designed to anneal to nucleotides 175 to 194 downstream of the translation start codon (nucleotide 1) of the *FMRFa* gene. Complementary oligos were annealed and cloned into the pCFD3-dU6:3gRNA vector (Addgene #49410). The vector was introduced into *vermillion* (*v*) strain and crossed with *Act5C-Cas9* flies (Bloomington #54590). Approximately 10 progenies were collected and established as independent lines. Each fly within the established lines were sequenced. The deletion in *ΔFMRFa* removes almost 900 nucleotides (175 to 1063) encoding all 14 *FMRFa* peptides. Glu700KDEL was a gift from Dr. Wolf Frommer (Addgene plasmid #17867).

### Immunostaining

Fly brains and thoracic ganglions were dissected in 1xPBS and then fixed in 4% paraformaldehyde with 0.2% Triton X-100 for 30 min at room temperature. Fixed brains were washed three times with PBS containing 0.2% Triton X-100. The samples were incubated in blocking buffer (5% heat-inactivated goat serum) with 0.2% Triton X-100 for 30 min at room temperature. Then, samples were incubated with 1/1000 diluted primary antibodies for 1-2 days at 4 °C. Samples were washed three times with PBS with 0.2% Triton X-100 and incubated with 1/1000 diluted secondary antibodies for 1-2 days at 4 °C. Samples were washed three times with PBS with 0.2% Triton X-100 and mounting solution (Vectashield) was added. Primary antibodies used in this study include chicken anti-GFP (Thermo Fisher Scientific), rabbit anti-mCherry (BioVision), rabbit anti-NPF (RayBiotech), mouse anti- Orcokinin-A (gift from Dr. Mizoguchi), rabbit anti-IPNa (gift from Dr. Thor), rabbit anti-Pdf (gift from Dr. Hardin), rabbit anti-FMRFa (RayBiotech). Secondary antibodies were anti-chicken Alexa488 (Thermo Fisher Scientific), anti-rabbit Alexa555 (Thermo Fisher Scientific) and anti-mouse Cy3 (The Jackson laboratory).

### *In vivo* Glucose imaging

Glucose imaging of FMRF neuron in the thoracic ganglion was conducted analogously as reported for NPF brain neurons, with the following modifications: the thoracic ganglion was extracted from the thorax while liquid silicone was applied over the entire preparation to prevent desiccation and preserve intracellular homeostatic conditions. Imaging was performed with a Nikon Eclipse Ti inverted microscope using 20x water objective, a dichroic filter (Nikon; 89002), excitation and emission filter wheels with four Chroma filters ET430/24x (#234435), ET500/20x (#235394), ET470/24m (#234331) and ET535/30m (Chroma #239226).

The light source was a Lumen 200 lamp (Prior Scientific Inc). Data acquisition was performed with NIS-Elements software (Nikon). For each data point, three images were sequentially obtained to calculate FRET (420-445 nm for CFP excitation and 458-482 nm for CFP emission with 400 ms exposure (Dd), 420-445 nm for CFP excitation and 520-550 nm for YFP emission with 100 ms exposure (Da), 491-508 nm for YFP excitation and 520-550 nm for YFP emission with 100 ms exposure (Aa)). Dd and Aa were used to calculate false FRET signals generated by CFP and YFP molecules alone, respectively. Spillover factors for CFP (0.290) and YFP (0.095) were experimentally established (Miyamoto and Amrein, 2019). For data analysis, adjacent area of cell bodies was subtracted as a background. FRET efficiency was calculated using a following formula; (Da-0.29xDd-0.095xAa)/Dd.

### Golgi size measurement

The Golgi apparatus was visualized using *UAS-ManII-eGFP* expressed under the control of respective GAL4 drivers. Golgi and NP staining was performed as described in Immunostaining except for using 4% paraformaldehyde with 0.01% Triton X-100 for fixation. All images were taken using a Nikon A1 confocal microscope with 60 x oil lens. Golgi and NP images were stacked (0.5 µm section intervals) and analyzed using ImageJ. Background was subtracted by thresholding using MaxEntropy for Golgi, Huang or Mean for NPs, and remained pixel numbers were counted. Golgi area was determined as the fraction of the area positive for NP, which served as a proxy for the cytosol.

### Metabolic Analyses

Glucose/trehalose measurements: For glucose and trehalose measurements, a single male fly was homogenized in 100 μl of water. After heat-inactivation at 95 °C for 5 min, non-soluble debris was removed by centrifugation for 10 minutes at 14000 rpm and soluble fraction was transferred to a new Eppendorf tube. 5 μl of the soluble fraction was used for determination of glucose and trehalose content, respectively. Glucose was determined by adding 100 μl of Infinity^TM^ Glucose Hexokinase Reagent (Thermo Scientific) to 5 μl of the soluble fraction for 10 minutes at 37 °C. Absorbance was measured at 340 nm using a Genesys 20 spectrophotometer (Thermo Scientific) and quantified against a standard glucose curve. For trehalose measurement, 5 μl of porcine kidney trehalase (2.3 units/mg; T8778-1UN, Sigma) was added to 5 μl of the soluble fraction for 30 minutes at 37 °C, followed the addition of 100 μl of the Infinity^TM^ Glucose Hexokinase Reagent. Absorbance was measured as for glucose, and final trehalose amount was determined by deducting glucose content of the same fly sample. Fed flies were taken straight from standard food, while starved flies are flies kept on 1% agarose for 24 hours prior measurement.

Glycogen measurement: Adult males were anesthetized using CO2. The abdomen and head were removed, the thorax separated into the dorsal (flight muscle) and the ventral halves (jump muscle). Jump muscles from three flies were placed into an Eppendorf tube containing 50 µl of 0.2 % NP-40 and stored at - 80 °C, until further processing. For measurements, samples were thawed, homogenized using a plastic pestle, and non-soluble debris was separated by centrifugation at 4 °C for 20 minutes at 14000 rpm. 5 µl of the supernatant was incubated with 5 µg of amyloglucosidase (Sigma- Aldrich A1602) with 50 mM Na acetate for 30 minutes at 37 °C. The aliquot was mixed with 100 µl of Infinity^TM^ Glucose Hexokinase Reagent and incubated for 10 minutes at 37 °C, before absorbance was measured at 340 nm. Glycogen amount was obtained by subtraction of free glucose which was determined by measuring 340 nm absorbance of 5 µl of the same sample in the absence of amyloglucosidase. Samples were normalized to total amount of protein, which was determined using 20 µl of supernatant, to which with 20 µl water was added. 2.5 µl of this was mixed with 100 µl of Bradford reagent (Bio-Rad #5000205) and absorbance was taken at 595 nm to calculate protein concentration against a standard curve.

### Hemolymph neuropeptide measurements

Hemolymph extraction was performed as described by Davoodi and colleagues (Davoodi et al., 2019). Briefly, abdomens of ten flies were punctured at the posterior end and transferred in an iced Eppendorf tube containing 60 µl of PBS with 0.2 % Tween20. Flies were submerged in the buffer by spinning samples for 1 minute at 3000 × g in a microcentrifuge, followed by vortexing at speed 10 (Fisher Scientific Vortex mixer) for 10 minutes. Debris was removed by centrifugation for 1 minute at 3000 × g, and the supernatant (hemolymph) was transferred to a new tube.

ELISA was performed in triplicates using the ANP ELISA kit (Abcam #ab108797). Briefly, 50 µl of samples and standard titration in the range from 0.008 ng/mL to 2 ng/mL were added to each precoated well. The plate was fully sealed and incubated for 2-3 hours, followed by 4-5 washes with 200 µl of wash buffer. 50 µl of Biotinylated ANP antibody was added and incubated for 2 hours at room temperature. The wells were washed three times and 50 µl of Streptavidin-Peroxidase conjugate was added. After 30 minutes, the wells were washed once. 50 µl of chromogen substrate was added and incubated for 20 minutes or until the desired blue color was achieved, after which the reaction was terminated by adding 50 µl of the stop solution. Absorbance was measured immediately at 450nm using a microplate reader (BioTek Instruments Inc).

Measurement of Ilp2HF in the hemolymph was performed using the *Ilp2^1^ gd2HF* flies as described previously (Park et al 2014). Plates (Nunc Immuno clear modules; Thermo Fisher Scientific 468667) were coated with anti-Flag antibody (Sigma-Aldrich F1804) diluted in 0.2 M sodium carbonate/bicarbonate buffer (pH 9.4) to a final concentration of 2.5µg/mL. Plates were incubated overnight at 4 °C, washed twice using PBS containing 0.2% Tween 20. The wells were coated with 350µL PBS containing 2% BSA overnight, washed three times with PBS containing 0.2% Tween 20. The hemolymph was extracted in the same way as Anf-GFP, described above. 50µL of hemolymph containing solution or FLAG(GS)HA peptide standard (DYKDDDDKGGGGSYPYDVPDYAamide, life-Tein LLC) was mixed with 50µl of PBS containing 1% Tween20 and 5ng/mL anti-HA-Peroxidase 3F10 antibody (Roche #12013819001) and transferred to the wells. Plates were sealed and incubated overnight at 4 °C. Samples were removed and wells were washed six times with PBS with 0.2% Tween20. 100µL of 1-Step Ultra TMB ELISA substrate (Thermo Fisher Scientific #34028) was added to each well, and plates were gently agitated for 30 minutes at room temperature. The reaction was stopped by adding 100µL of 2M sulfuric acid, and absorbance was measured immediately at 450nm.

### Climbing assay

The climbing assay was performed as reported by Ali and colleagues (Ali et al., 2011). Approximately 15-20 male flies were placed in a culture plastic vial containing regular food (fed conditions) or plain agarose (starved conditions) 24 hours prior to testing. Flies were transferred into an empty vial with a line marker 8 cm from the bottom, and tested by tapping them down gently to the bottom. The number of flies climbing up to marked within 10 seconds were counted.

## REFERENCES

1. Ali, Y.O., Escala, W., Ruan, K., Zhai, R.G., 2011. Measurement of the Relations between Chromosomes and Behavior. Journal of visualized experiments: JoVE. 10.3791/2504

2. Baines, R.A., Uhler, J.P., Thompson, A., Sweeney, S.T., Bate, M., 2001. Altered electrical properties in Drosophila neurons developing without synaptic transmission. J Neurosci 21, 1523–1531. 10.1523/JNEUROSCI.21-05-01523.2001

3. Banka, S., Chervinsky, E., Newman, W.G., Crow, Y.J., Yeganeh, S., Yacobovich, J., Donnai, D., Shalev, S., 2011. Further delineation of the phenotype of severe congenital neutropenia type 4 due to mutations in G6PC3. Eur J Hum Genet 19, 18–22. 10.1038/ejhg.2010.136

4. Bosma, K.J., Rahim, M., Singh, K., Goleva, S.B., Wall, M.L., Xia, J., Syring, K.E., Oeser, J.K., Poffenberger, G., McGuinness, O.P., Means, A.L., Powers, A.C., Li, W.-H., Davis, L.K., Young, J.D., O’Brien, R.M., 2020. Pancreatic islet beta cell-specific deletion of G6pc2 reduces fasting blood glucose. J Mol Endocrinol 64, 235–248. 10.1530/JME-20-0031

5. Brussaard, A.B., Kits, K.S., Ter Maat, A., Van Minnen, J., Moed, P.J., 1988. Dual inhibitory action of FMRFamide on neurosecretory cells controlling egg laying behavior in the pond snail. Brain Res 447, 35–51. 10.1016/0006-8993(88)90963-8

6. Bryantsev, A.L., Baker, P.W., Lovato, T.L., Jaramillo, M.S., Cripps, R.M., 2012. Differential requirements for Myocyte Enhancer Factor-2 during adult myogenesis in Drosophila. Dev Biol 361, 191–207. 10.1016/j.ydbio.2011.09.031

7. Cheung, Y.Y., Kim, S.Y., Yiu, W.H., Pan, C.-J., Jun, H.-S., Ruef, R.A., Lee, E.J., Westphal, H., Mansfield, B.C., Chou, J.Y., 2007. Impaired neutrophil activity and increased susceptibility to bacterial infection in mice lacking glucose-6-phosphatase-beta. J Clin Invest 117, 784–793. 10.1172/JCI30443

8. Chung, B.Y., Ro, J., Hutter, S.A., Miller, K.M., Guduguntla, L.S., Kondo, S., Pletcher, S.D., 2017. Drosophila Neuropeptide F Signaling Independently Regulates Feeding and Sleep-Wake Behavior. Cell Rep 19, 2441–2450. 10.1016/j.celrep.2017.05.085

9. Davoodi, S., Galenza, A., Panteluk, A., Deshpande, R., Ferguson, M., Grewal, S., Foley, E., 2019. The Immune Deficiency Pathway Regulates Metabolic Homeostasis in Drosophila. J Immunol 202, 2747–2759. 10.4049/jimmunol.1801632

10. Demontis, F., Piccirillo, R., Goldberg, A.L., Perrimon, N., 2013. The influence of skeletal muscle on systemic aging and lifespan. Aging Cell 12, 943–949. 10.1111/acel.12126

11. Deng, B., Li, Q., Liu, X., Cao, Y., Li, B., Qian, Y., Xu, R., Mao, R., Zhou, E., Zhang, W., Huang, J., Rao, Y., 2019. Chemoconnectomics: Mapping Chemical Transmission in Drosophila. Neuron 101, 876–893.e4. 10.1016/j.neuron.2019.01.045

12. Fisher, T., Lin, C.H., Kaczmarek, L.K., 1993. The peptide FMRFa terminates a discharge in Aplysia bag cell neurons by modulating calcium, potassium, and chloride conductances. J Neurophysiol 69, 2164–2173. 10.1152/jn.1993.69.6.2164

13. Fujii, S., Ahn, J.-E., Jagge, C., Shetty, V., Janes, C., Mohanty, A., Slotman, M., Adelman, Z.N., Amrein, H., 2023. RNA Taste is Conserved in Dipteran Insects. J Nutr S0022–3166(23)35477–4. 10.1016/j.tjnut.2023.03.010

14. Gauthier, S.A., Hewes, R.S., 2006. Transcriptional regulation of neuropeptide and peptide hormone expression by the Drosophila dimmed and cryptocephal genes. J Exp Biol 209, 1803–1815. 10.1242/jeb.02202

15. Gomarasca, M., Banfi, G., Lombardi, G., 2020. Myokines: The endocrine coupling of skeletal muscle and bone. Adv Clin Chem 94, 155–218. 10.1016/bs.acc.2019.07.010

16. Hardt, B., Völker, C., Mundt, S., Salska-Navarro, M., Hauptmann, M., Bause, E., 2005. Human endo-alpha1,2-mannosidase is a Golgi-resident type II membrane protein. Biochimie 87, 169–179. 10.1016/j.biochi.2004.11.004

17. Hayee, B., Antonopoulos, A., Murphy, E.J., Rahman, F.Z., Sewell, G., Smith, B.N., McCartney, S., Furman, M., Hall, G., Bloom, S.L., Haslam, S.M., Morris, H.R., Boztug, K., Klein, C., Winchester, B., Pick, E., Linch, D.C., Gale, R.E., Smith, A.M., Dell, A., Segal, A.W., 2011. G6PC3 mutations are associated with a major defect of glycosylation: a novel mechanism for neutrophil dysfunction. Glycobiology 21, 914–924. 10.1093/glycob/cwr023

18. Herzog, H., Darby, K., Ball, H., Hort, Y., Beck-Sickinger, A., Shine, J., 1997. Overlapping gene structure of the human neuropeptide Y receptor subtypes Y1 and Y5 suggests coordinate transcriptional regulation. Genomics 41, 315–319. 10.1006/geno.1997.4684

19. Hewes, R.S., Park, D., Gauthier, S.A., Schaefer, A.M., Taghert, P.H., 2003. The bHLH protein Dimmed controls neuroendocrine cell differentiation in Drosophila. Development 130, 1771–1781. 10.1242/dev.00404

20. Higuchi, H., Hasegawa, A., Yamaguchi, T., 2005. Transcriptional regulation of neuronal genes and its effect on neural functions: transcriptional regulation of neuropeptide Y gene by leptin and its effect on feeding. J Pharmacol Sci 98, 225–231. 10.1254/jphs.fmj05001x6

21. Hook, V., Funkelstein, L., Lu, D., Bark, S., Wegrzyn, J., Hwang, S.-R., 2008. Proteases for processing proneuropeptides into peptide neurotransmitters and hormones. Annu Rev Pharmacol Toxicol 48, 393–423. 10.1146/annurev.pharmtox.48.113006.094812

22. Husain, Q.M., Ewer, J., 2004. Use of targetable gfp-tagged neuropeptide for visualizing neuropeptide release following execution of a behavior. J Neurobiol 59, 181–191. 10.1002/neu.10309

23. Kim, S.K., Tsao, D.D., Suh, G.S.B., Miguel-Aliaga, I., 2021. Discovering signaling mechanisms governing metabolism and metabolic diseases with Drosophila. Cell Metab 33, 1279– 1292. 10.1016/j.cmet.2021.05.018

24. Kneen, M., Farinas, J., Li, Y., Verkman, A.S., 1998. Green fluorescent protein as a noninvasive intracellular pH indicator. Biophys J 74, 1591–1599. 10.1016/S0006-3495(98)77870-1

25. Kondylis, V., Rabouille, C., 2009. The Golgi apparatus: lessons from Drosophila. FEBS Lett 583, 3827–3838. 10.1016/j.febslet.2009.09.048

26. Kovac, S., Shulkes, A., Baldwin, G.S., 2009. Peptide processing and biology in human disease. Curr Opin Endocrinol Diabetes Obes 16, 79–85. 10.1097/MED.0b013e3283202555

27. Landayan, D., Wang, B.P., Zhou, J., Wolf, F.W., 2021. Thirst interneurons that promote water seeking and limit feeding behavior in Drosophila. Elife 10. 10.7554/eLife.66286

28. Lear, B.C., Zhang, L., Allada, R., 2009. The neuropeptide PDF acts directly on evening pacemaker neurons to regulate multiple features of circadian behavior. PLoS Biol 7, e1000154. 10.1371/journal.pbio.1000154

29. Lee, K.-S., You, K.-H., Choo, J.-K., Han, Y.-M., Yu, K., 2004. Drosophila short neuropeptide F regulates food intake and body size. J Biol Chem 279, 50781–50789. 10.1074/jbc.M407842200

30. Lenz, O., Xiong, J., Nelson, M.D., Raizen, D.M., Williams, J.A., 2015. FMRFamide signaling promotes stress-induced sleep in Drosophila. Brain Behav Immun 47, 141–148. 10.1016/j.bbi.2014.12.028

31. Liu, W., Ganguly, A., Huang, J., Wang, Y., Ni, J.D., Gurav, A.S., Aguilar, M.A., Montell, C., 2019. Neuropeptide F regulates courtship in Drosophila through a male-specific neuronal circuit. Elife 8. 10.7554/eLife.49574

32. Lu, T.-C., Brbić, M., Park, Y.-J., Jackson, T., Chen, J., Kolluru, S.S., Qi, Y., Katheder, N.S., Cai, X.T., Lee, S., Chen, Y.-C., Auld, N., Liang, C.-Y., Ding, S.H., Welsch, D., D’Souza, S., Pisco, A.O., Jones, R.C., Leskovec, J., Lai, E.C., Bellen, H.J., Luo, L., Jasper, H., Quake, S.R., Li, H., 2023. Aging Fly Cell Atlas identifies exhaustive aging features at cellular resolution. Science 380, eadg0934. 10.1126/science.adg0934

33. Luan, H., Peabody, N.C., Vinson, C.R., White, B.H., 2006. Refined spatial manipulation of neuronal function by combinatorial restriction of transgene expression. Neuron 52, 425–436. 10.1016/j.neuron.2006.08.028

34. Lundquist, T., Nässel, D.R., 1990. Substance P-, FMRFamide-, and gastrin/cholecystokinin-like immunoreactive neurons in the thoraco-abdominal ganglia of the flies Drosophila and Calliphora. J Comp Neurol 294, 161–178. 10.1002/cne.902940202

35. Matsuda, H., Yamada, T., Yoshida, M., Nishimura, T., 2015. Flies without trehalose. J Biol Chem 290, 1244–1255. 10.1074/jbc.M114.619411

36. Miyamoto, T., Amrein, H., 2019. Neuronal Gluconeogenesis Regulates Systemic Glucose Homeostasis in Drosophila melanogaster. Curr Biol 29, 1263–1272.e5. 10.1016/j.cub.2019.02.053

37. Miyamoto, T., Amrein, H., 2017. Gluconeogenesis: An ancient biochemical pathway with a new twist. Fly (Austin) 11, 218–223. 10.1080/19336934.2017.1283081

38. Miyamoto, T., Amrein, H., 2014. Diverse roles for the Drosophila fructose sensor Gr43a. Fly (Austin) 8, 19–25. 10.4161/fly.27241

39. Miyamoto, T., Slone, J., Song, X., Amrein, H., 2012. A fructose receptor functions as a nutrient sensor in the Drosophila brain. Cell 151, 1113–1125. 10.1016/j.cell.2012.10.024

40. Nässel, D.R., Zandawala, M., 2019. Recent advances in neuropeptide signaling in Drosophila, from genes to physiology and behavior. Prog Neurobiol 179, 101607. 10.1016/j.pneurobio.2019.02.003

41. Oh, Y., Lai, J.S.-Y., Mills, H.J., Erdjument-Bromage, H., Giammarinaro, B., Saadipour, K., Wang, J.G., Abu, F., Neubert, T.A., Suh, G.S.B., 2019. A glucose-sensing neuron pair regulates insulin and glucagon in Drosophila. Nature 574, 559–564. 10.1038/s41586-019-1675-4

42. Paradis, S., Sweeney, S.T., Davis, G.W., 2001. Homeostatic control of presynaptic release is triggered by postsynaptic membrane depolarization. Neuron 30, 737–749. 10.1016/s0896-6273(01)00326-9

43. Parisky, K.M., Agosto, J., Pulver, S.R., Shang, Y., Kuklin, E., Hodge, J.J.L., Kang, K., Liu, X., Garrity, P.A., Rosbash, M., Griffith, L.C., 2008. PDF cells are a GABA-responsive wake-promoting component of the Drosophila sleep circuit. Neuron 60, 672–682. 10.1016/j.neuron.2008.10.042

44. Park, S., Alfa, R.W., Topper, S.M., Kim, G.E.S., Kockel, L., Kim, S.K., 2014. A genetic strategy to measure circulating Drosophila insulin reveals genes regulating insulin production and secretion. PLoS Genet 10, e1004555. 10.1371/journal.pgen.1004555

45. Pauls, D., Chen, J., Reiher, W., Vanselow, J.T., Schlosser, A., Kahnt, J., Wegener, C., 2014. Peptidomics and processing of regulatory peptides in the fruit fly Drosophila. EuPA Open Proteomics 3, 114–127.

46. Pfeiffer, B.D., Ngo, T.-T.B., Hibbard, K.L., Murphy, C., Jenett, A., Truman, J.W., Rubin, G.M., 2010. Refinement of tools for targeted gene expression in Drosophila. Genetics 186, 735–755. 10.1534/genetics.110.119917

47. Predel, R., Wegener, C., Russell, W.K., Tichy, S.E., Russell, D.H., Nachman, R.J., 2004. Peptidomics of CNS-associated neurohemal systems of adult Drosophila melanogaster: a mass spectrometric survey of peptides from individual flies. J Comp Neurol 474, 379–392. 10.1002/cne.20145

48. Rai, M., Demontis, F., 2016. Systemic Nutrient and Stress Signaling via Myokines and Myometabolites. Annu Rev Physiol 78, 85–107. 10.1146/annurev-physiol-021115-105305

49. Ravi, P., Trivedi, D., Hasan, G., 2018. FMRFa receptor stimulated Ca2+ signals alter the activity of flight modulating central dopaminergic neurons in Drosophila melanogaster. PLoS Genet 14, e1007459. 10.1371/journal.pgen.1007459

50. Renn, S.C., Park, J.H., Rosbash, M., Hall, J.C., Taghert, P.H., 1999. A pdf neuropeptide gene mutation and ablation of PDF neurons each cause severe abnormalities of behavioral circadian rhythms in Drosophila. Cell 99, 791–802. 10.1016/s0092-8674(00)81676-1

51. Rholam, M., Fahy, C., 2009. Processing of peptide and hormone precursors at the dibasic cleavage sites. Cell Mol Life Sci 66, 2075–2091. 10.1007/s00018-009-0007-5

52. Schneider, L.E., Roberts, M.S., Taghert, P.H., 1993a. Cell type-specific transcriptional regulation of the Drosophila FMRFamide neuropeptide gene. Neuron 10, 279–291. 10.1016/0896-6273(93)90318-l

53. Schneider, L.E., Sun, E.T., Garland, D.J., Taghert, P.H., 1993b. An immunocytochemical study of the FMRFamide neuropeptide gene products in Drosophila. J Comp Neurol 337, 446–460. 10.1002/cne.903370308

54. Shao, L., Saver, M., Chung, P., Ren, Q., Lee, T., Kent, C.F., Heberlein, U., 2017. Dissection of the Drosophila neuropeptide F circuit using a high-throughput two-choice assay. Proc Natl Acad Sci U S A 114, E8091–E8099. 10.1073/pnas.1710552114

55. Silva, V., Palacios-Muñoz, A., Volonté, M., Frenkel, L., Ewer, J., Ons, S., 2021. Orcokinin neuropeptides regulate reproduction in the fruit fly, Drosophila melanogaster. Insect Biochem Mol Biol 139, 103676. 10.1016/j.ibmb.2021.103676

56. Song, T., Qin, W., Lai, Z., Li, H., Li, D., Wang, B., Deng, W., Wang, T., Wang, L., Huang, R., 2023. Dietary cysteine drives body fat loss via FMRFamide signaling in Drosophila and mouse. Cell Res 33, 434–447. 10.1038/s41422-023-00800-8

57. Taghert, P.H., Hewes, R.S., Park, J.H., O’Brien, M.A., Han, M., Peck, M.E., 2001. Multiple amidated neuropeptides are required for normal circadian locomotor rhythms in Drosophila. J Neurosci 21, 6673–6686. 10.1523/JNEUROSCI.21-17-06673.2001

58. Thompson, S.N., 2003. Trehalose – The Insect ‘Blood’ Sugar. Advances in Insect Physiology 31, 205–285. 10.1016/S0065-2806(03)31004-5

59. van den Pol, A.N., 2012. Neuropeptide transmission in brain circuits. Neuron 76, 98–115. 10.1016/j.neuron.2012.09.014

60. van Schaftingen, E., Gerin, I., 2002. The glucose-6-phosphatase system. Biochem J 362, 513–532. 10.1042/0264-6021:3620513

61. Veiga-da-Cunha, M., Chevalier, N., Stephenne, X., Defour, J.-P., Paczia, N., Ferster, A., Achouri, Y., Dewulf, J.P., Linster, C.L., Bommer, G.T., Van Schaftingen, E., 2019. Failure to eliminate a phosphorylated glucose analog leads to neutropenia in patients with G6PT and G6PC3 deficiency. Proc Natl Acad Sci U S A 116, 1241–1250. 10.1073/pnas.1816143116

62. Verleyen, P., Baggerman, G., Wiehart, U., Schoeters, E., Van Lommel, A., De Loof, A., Schoofs, L., 2004. Expression of a novel neuropeptide, NVGTLARDFQLPIPNamide, in the larval and adult brain of Drosophila melanogaster. J Neurochem 88, 311–319. 10.1046/j.1471-4159.2003.02161.x

63. Volkenhoff, A., Weiler, A., Letzel, M., Stehling, M., Klämbt, C., Schirmeier, S., 2015. Glial Glycolysis Is Essential for Neuronal Survival in Drosophila. Cell Metab 22, 437–447. 10.1016/j.cmet.2015.07.006

64. Wu, Q., Wen, T., Lee, G., Park, J.H., Cai, H.N., Shen, P., 2003. Developmental control of foraging and social behavior by the Drosophila neuropeptide Y-like system. Neuron 39, 147–161. 10.1016/s0896-6273(03)00396-9

65. Zhao, X., Karpac, J., 2017. Muscle Directs Diurnal Energy Homeostasis through a Myokine-Dependent Hormone Module in Drosophila. Curr Biol 27, 1941–1955.e6. 10.1016/j.cub.2017.06.004

66. Zhao, X., Li, X., Shi, X., Karpac, J., 2020. Diet-MEF2 interactions shape lipid droplet diversification in muscle to influence Drosophila lifespan. Aging Cell 19, e13172. 10.1111/acel.13172

67. Zhou, W., Chang, J., Wang, X., Savelieff, M.G., Zhao, Y., Ke, S., Ye, B., 2014. GM130 is required for compartmental organization of dendritic golgi outposts. Curr Biol 24, 1227–1233. 10.1016/j.cub.2014.04.008

