## supplemental data for "*Drosophila* Neuronal Glucose 6 Phosphatase is a Modulator of Neuropeptide Release that Regulates Muscle Glycogen Stores via FMRFamide Signaling"

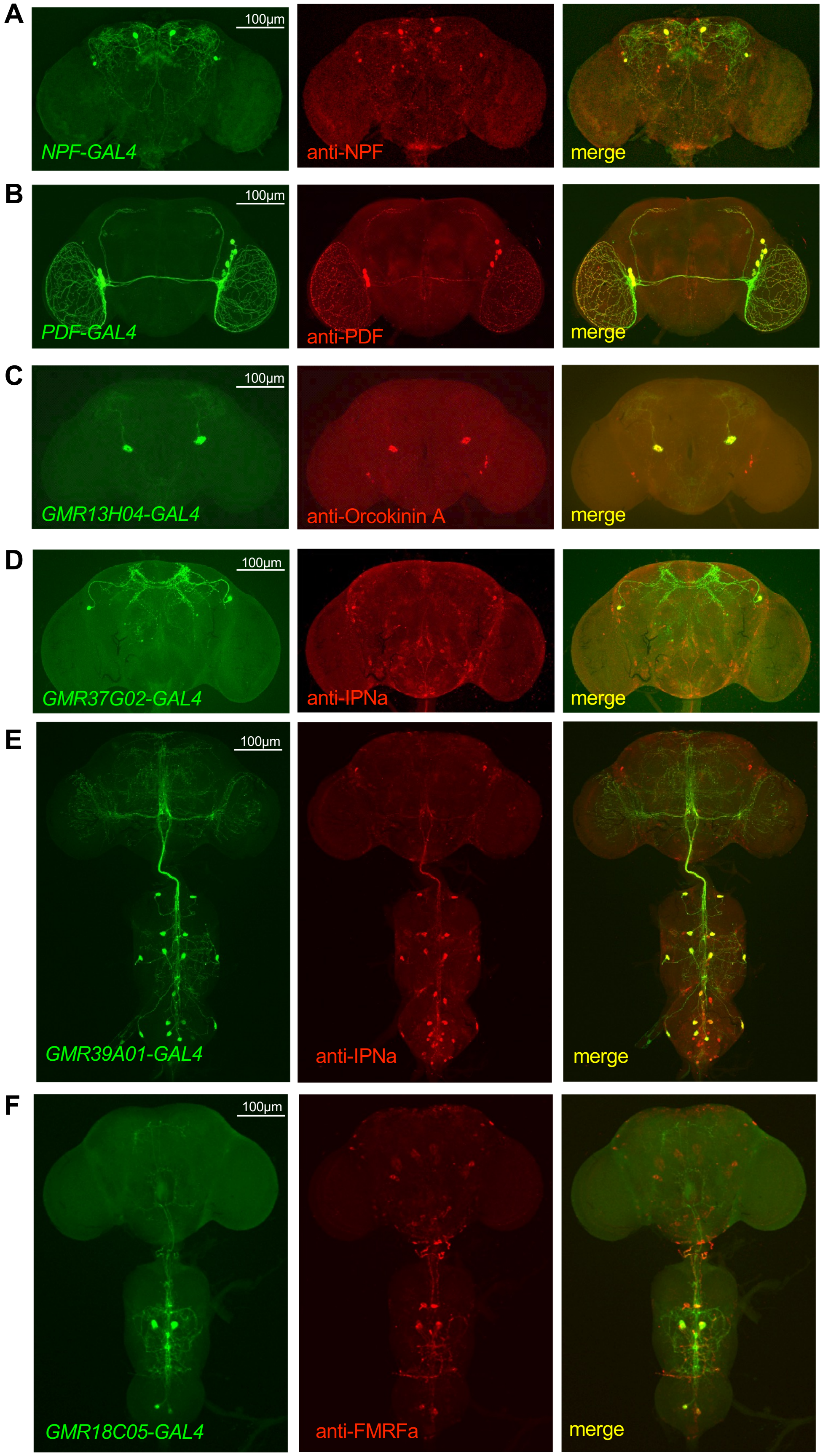


**Supplementary Figure 1**

**Supplementary Figure 1: Identification and Expression of Gal4 lines labeling specific peptidergic neurons**

**A-B)** Brains of flies expressing *UAS-mCD8GFP* driven by *NPF-GAL4* (Wu et al., 2003) and *PDF-GAL4* (Renn et al., 1999) were double-stained with antibodies against GFP and NPF **(A)** and PDF **(B)** respectively, revealing co-expression in all GAL4 expressing neurons.

**C-F)** Neuropeptide specific *FlyLight GAL4* lines *GMR13H04*, *GMR37G02*, *GMR39A01* and *GMR18C05* were identified by comparing expression profiles with antibody staining data (Nässel and Zandawala, 2019, and references therein). CNS of these lines in the presence of a *UAS-mCD8GFP* reporter were double-stained with anti-GFP and NP-specific antibodies, which revealed that the respective *GAL4* drivers were expressed in large Orcokinin A **(C)**, IPNa **(D and E)** and FMRFamides **(F)** neurons, respectively.

A

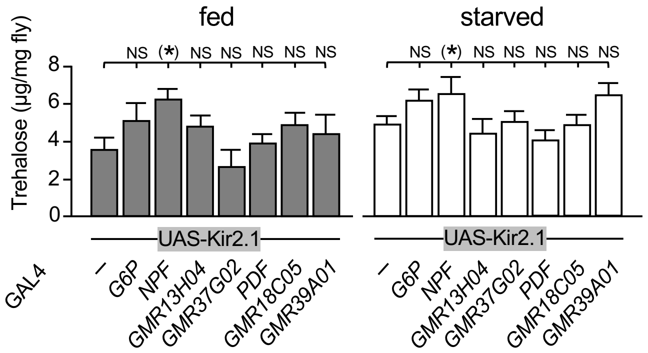


**Supplementary Figure 2. Whole body trehalose does not significantly differ when subsets most subsets of *G6P* neurons are inactivated by Kir2.1.** Trehalose amounts of experimental flies was compared to that of flies containing only the *UAS-Kir2.1* reporter. Trehalose amount was increased when *NPF* neurons were inactivated. Asterisks indicate *P<0.05 by Kruskal-Wallis with Dunn's post-hoc test; n=8-10. Error bars represent standard error.

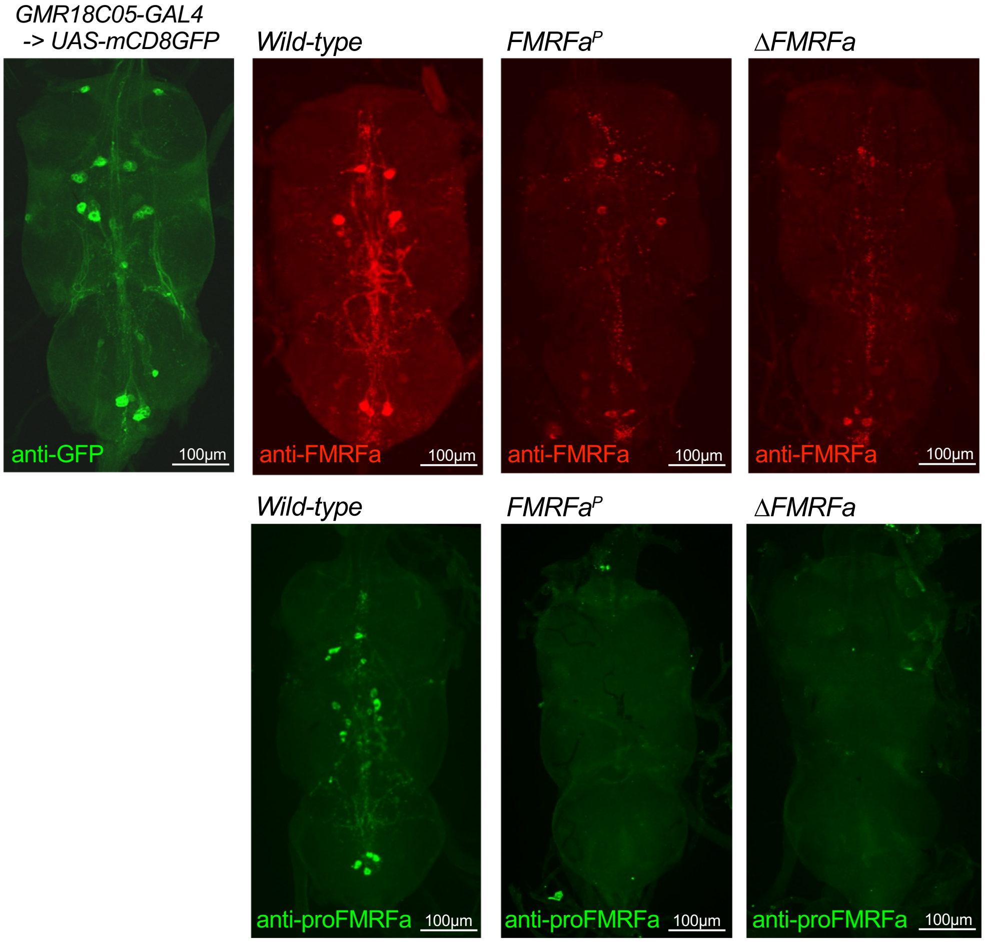


**Supplementary Figure 3. Thoracic ganglia expression of FMRFamides in the *wild-type*, *FMRFa[P]* and Δ*FMRFa* mutant flies.** Immunostaining was performed using anti-FMRFa antibody (raised against a FMRF peptide) and anti-proFMRFa antibodies (raised against the C-terminal domain of FMRFa pre-protein). Very weak Anti-FMRFa antibody staining can be seen in *FMRFa*$^{P}$ and Δ*FMRFa* mutants, derived possibly from low amounts (*FMRFa*$^{P}$) and/or cross-reactivity to related tetrapeptides (FLRF in Dromyosuppressin, HMRF in Drosulfakinin), Importantly, no staining was observed in mutants using anti-proFMRFa antibody.

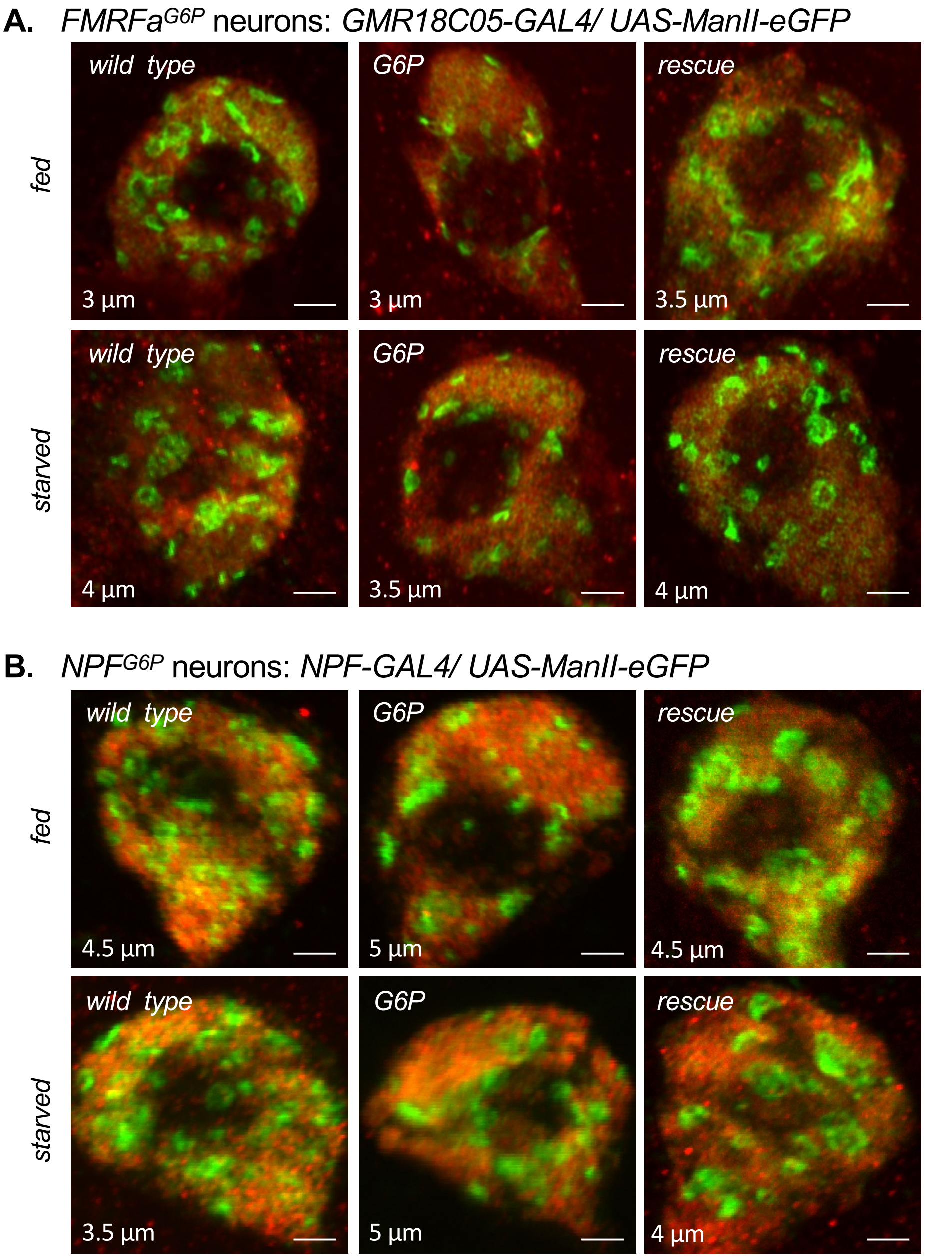


**Supplementary Figure 4**

**Supplementary Figure 4. Enlarged images of Golgi apparatus in the *FMRFa*$^{G6P}$ and *NPF*$^{G6P}$ neurons.** Images (same as shown in Figure 4) represent maximum intensity of stacks ranging between 3 µm to 5 µm (indicated at the bottom left) of 0.5 µm thick sections. All genotypes include indicated driver. Genotypes: *G6P*$^{+}$*/G6P*$^{+}$ (*wild type*), *G6P*$^{MIC}$*/G6P*$^{MIC}$ (G6P) and *G6P*$^{MIC}$*/G6P*$^{MIC}$*; UAS-G6P* (rescue).

**A)** Golgi was visualized in *FMRFa*$^{G6P}$ neurons using *UAS-ManII-eGFP* expressed under the control of *GMR18C05-GAL4* (present in all flies). Scale bars are 4 µm.

**B)** Golgi was visualized in *NPF*$^{G6P}$ neurons using *UAS-ManII-eGFP* expressed under the control of *NPF-GAL4* (present in all flies). Scale bars are 4 µm.

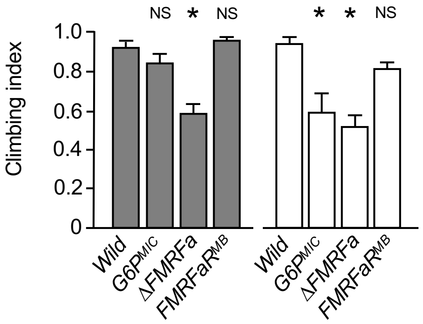


**Supplementary Figure 5. *G6P*$^{MIC}$ and ∆*FMRFa* mutant, but not *FMRFaR* mutant flies show reduced climbing performance.** The climbing assay was performed as described in Ali and colleagues (Ali et al., 2011) using fed and 24 hours food-deprived flies. ∆*FMRFa* mutant (fed and starved) and *G6P*$^{MIC}$ mutant flies (starved) show significantly diminished climbing performance compared to wild type and *FMRFaR* mutant flies. Asterisk indicates *P<0.05 using one-way ANOVA with Tukey post hoc test. N=7. Error bars represent standard error.

**Fly strains used in this study (from left to right)**

Figure 1: *w; G6P-GAL4; UAS-mCD8GFP*.

Figure 2: (A) *yw; UAS-Kir2.1*, *yw; G6P-GAL4/UAS-Kir2.1*, *yw; NPF-GAL4/UAS-Kir2.1*, *yw; UAS-Kir2.1; GMR13H04-GAL4*, *yw; UAS-Kir2.1; GMR37G02-GAL4*, *yw; UAS-Kir2.1; Pdf-GAL4*, *yw; UAS-Kir2.1; GMR18C05-GAL4*, *yw; UAS-Kir2.1; GMR39A01-GAL4*. (B) from left to right, *yw*, *yw;* $G6P^{MIC}/G6P^{MIC}$, *yw;* $G6P^{MIC}/G6P^{MIC}$*; GMR18C05-GAL4*, *yw;* $G6P^{MIC}/G6P^{MIC}$*; UAS-G6P*, *yw;* $G6P^{MIC}/G6P^{MIC}$*; GMR18C05-GAL4/UAS-G6P*. (D) from left to right, *yw*, *yw;* $G6P^{MIC}/G6P^{MIC}$, *yw;* $FMRFa^{P}/FMRFa^{P}$, *yw; ∆FMRFa/∆FMRFa*.

Figure 3: (A and D) *w; G6P-GAL4DBD FMRFa-p65AD/10xUAS-mCD8GFP*. (B and E) *w; GMR18C05-GAL4/UAS-mCD8GFP*.

Figure 4: *w; UAS-Glu700KDEL; GMR18C05-GAL4*, w;$G6P^{MIC}$ *UAS-Glu700KDEL/*$G6P^{MIC}$*; GMR18C05-GAL4*, *w;* $G6P^{MIC}$ *UAS-Glu700KDEL/*$G6P^{MIC}$*; GMR18C05-GAL4/UAS-G6P*.

Figure 5: (A) *w; UAS-ManII-eGFP; GMR18C05-GAL4*, w; $G6P^{MIC}$ *UAS-ManII-eGFP/*$G6P^{MIC}$*; GMR18C05-GAL4*, *w;* $G6P^{MIC}$ *UAS-ManII-eGFP/*$G6P^{MIC}$*; GMR18C05-GAL4/UAS-G6P*. (B) *w; NP-GAL4/UAS-ManII-eGFP*, w; $G6P^{MIC}$ *NP-GAL4/*$G6P^{MIC}$ *GMR18C05-GAL4*, w; $G6P^{MIC}$ *NP-GAL4/*$G6P^{MIC}$ *GMR18C05-GAL4; UAS-ManII-eGFP*.

Figure 6: (A) *UAS-AnfGFP; GMR18C05-GAL4*, *UAS-AnfGFP;* $G6P^{MIC}/G6P^{MIC}$*; GMR18C05-GAL4, UAS-AnfGFP;* $G6P^{MIC}/G6P^{MIC}$*; GMR18C05-GAL4/UAS-G6P.* (B) from left to right, *w;* $Ilp2^{1}$ *gd2HF/dilp2-GAL4, w;* $Ilp2^{1}$ *gd2HF/UAS-G6P, w;* $Ilp2^{1}$ *gd2HF/dilp2-GAL4 UAS-G6P.*

Figure 7: (A) *w;* $FMRFaR^{2A\text{-}GAL4}$*/20xUAS-6xGFP*. (B) *yw*, *yw;* $FMRFaR^{MB}$. (C) *yw*, *yw;* $FMRFaR^{MB}$, *Act79B-GAL4; ELAV-GAL80;* $FMRFaR^{MB}$, *yw;* $FMRFaR^{MB}$ *UAS-FMRFaR/*$FMRFaR^{MB}$*, Act79B-GAL4; ELAV-GAL80;* $FMRFaR^{MB}$ *UAS-FMRFaR/*$FMRFaR^{MB}$*, yw; ELAV-GAL80/CG-GAL4;* $FMRFaR^{MB}$ *UAS-FMRFaR/*$FMRFaR^{MB}$*.* (D) *yw*, *yw;* $G6P^{MIC}$, *yw; ∆FMRFa*.

*NPF-GAL4* (#25681), *GMR13H04-GAL4* (#48589), *GMR37G02-GAL4* (#49965), *GMR18C05-GAL4* (#47330), *GMR39A01-GAL4* (#45667), *CG-GAL4* (#7011), *UAS-Kir2.1* (#6596), UAS-ANF-GFP (#7001), *UAS-mCD8GFP* (#5130), *10xUAS-IVS-mCD8GFP* (#32185), *20xUAS-6xGFP* (#52262), *ELAV-GAL80*, *TI{2A-GAL4}*$FMRFaR^{2A\text{-}GAL4}$ (#84633), *G6P[MI12250]* (#57904), *FMRFa[KG01300]* (#13717), *FMRFaR[MB04659]* (#24212) and *UAS-ManII-eGFP* (#65248) strains were obtained from the Bloomington Stock Center (Indiana University). *Pdf-GAL4, UAS-FMRFaR*, *Act79B-GAL4* and $Ilp2^{1}$ *gd2HF* flies were obtained from Drs. Paul Hardin, Gaiti Hasan, Richard Cripps and Seung Kim, respectively. *G6P-GAL4*, *UAS-G6P*, *G6P-GAL4DBD*, *FMRFa-p65AD* and *∆FMRFa* were generated as described below (Molecular Biology section).

### REFERENCES:

Ali, Y.O., Escala, W., Ruan, K., Zhai, R.G., 2011. Measurement of the Relations between Chromosomes and Behavior. Journal of visualized experiments: JoVE. https://doi.org/10.3791/2504

Nässel, D.R., Zandawala, M., 2019. Recent advances in neuropeptide signaling in Drosophila, from genes to physiology and behavior. Prog Neurobiol 179, 101607. https://doi.org/10.1016/j.pneurobio.2019.02.003

Renn, S.C., Park, J.H., Rosbash, M., Hall, J.C., Taghert, P.H., 1999. A pdf neuropeptide gene mutation and ablation of PDF neurons each cause severe abnormalities of behavioral circadian rhythms in Drosophila. Cell 99, 791–802. https://doi.org/10.1016/s0092-8674(00)81676-1

Wu, Q., Wen, T., Lee, G., Park, J.H., Cai, H.N., Shen, P., 2003. Developmental control of foraging and social behavior by the Drosophila neuropeptide Y-like system. Neuron 39, 147–161. https://doi.org/10.1016/s0896-6273(03)00396-9
